# Structure and Evolution of Constitutive Bacterial Promoters

**DOI:** 10.1101/2020.05.19.104232

**Authors:** Mato Lagator, Srdjan Sarikas, Magdalena Steinrück, David Toledo-Aparicio, Jonathan P. Bollback, Gasper Tkacik, Calin C. Guet

## Abstract

Predicting gene expression levels from any DNA sequence is a major challenge in biology. Using libraries with >25,000 random mutants, we developed a biophysical model that accounts for major features of σ^70^-binding bacterial promoters to accurately predict constitutive gene expression levels of any sequence. We experimentally and theoretically estimated that 10-20% of random sequences lead to expression and 82% of non-expressing sequences are one point mutation away from a functional promoter. Generating expression from random sequences is pervasive, such that selection acts against σ^70^-RNA polymerase binding sites even within inter-genic, promoter-containing regions. The pervasiveness of σ^70^– binding sites, which arises from the structural features of promoters captured by our biophysical model, implies that their emergence is unlikely the limiting step in gene regulatory evolution.

## Main text

Gene expression is one of the most fundamental process of life, and tuning expression levels underpins complex biological function (*1*). The fundamental process in the expression of most bacterial genes is the recruitment of σ^70^-RNA Polymerase (RNAP) to a stretch of DNA – the promoter. The relationship between promoter sequence (the genotype) and gene expression levels (the phenotype) is central to understanding cellular function and the evolution of gene regulatory networks (*2*).

Attempts to understand genotype-phenotype relationship in promoters have adopted, broadly speaking, three approaches. (i) Bioinformatics identifies promoters based on sequence homology to the σ^70^-RNAP consensus site (*3*), which consists of -10 and -35 elements (TATAAT and TTGACA in *E.coli*, respectively) separated by a spacer with canonical length of 17bp, but does not predict gene expression from them (*4*). (ii) Machine learning accurately predicts gene expression patterns in cells (*5*), but lacks direct links to the underlying biological mechanisms, limiting insights into promoter structure and evolution (*6*). (iii) Biophysical models, the most successful of which predict gene expression levels based on the thermodynamic properties of σ^70^-RNAP binding at a promoter in equilibrium (*7*), do not generalize to random sequences (*8*). In sum, we lack a generalizable and predictive understanding of the relationship between promoter genotype and gene expression phenotype even for constitutive promoters, where σ^70^-RNAP binding solely determines gene expression levels (*9*).

The standard thermodynamic model (*7, 10*) considers the energy of binding between σ^70^-RNAP complex and DNA to identify, typically, the single strongest binding site, with the expression level proportional to the equilibrium occupancy of σ^70^-RNAP on that binding site (‘Standard’ model, Fig.1A). To predict gene expression levels from any random sequence, we developed a comprehensive and generalizable thermodynamic model (‘Extended’) that accounts for six essential *structural features of bacterial promoters* that are not present in the Standard model (Fig.1A,B).

**Figure 1.**
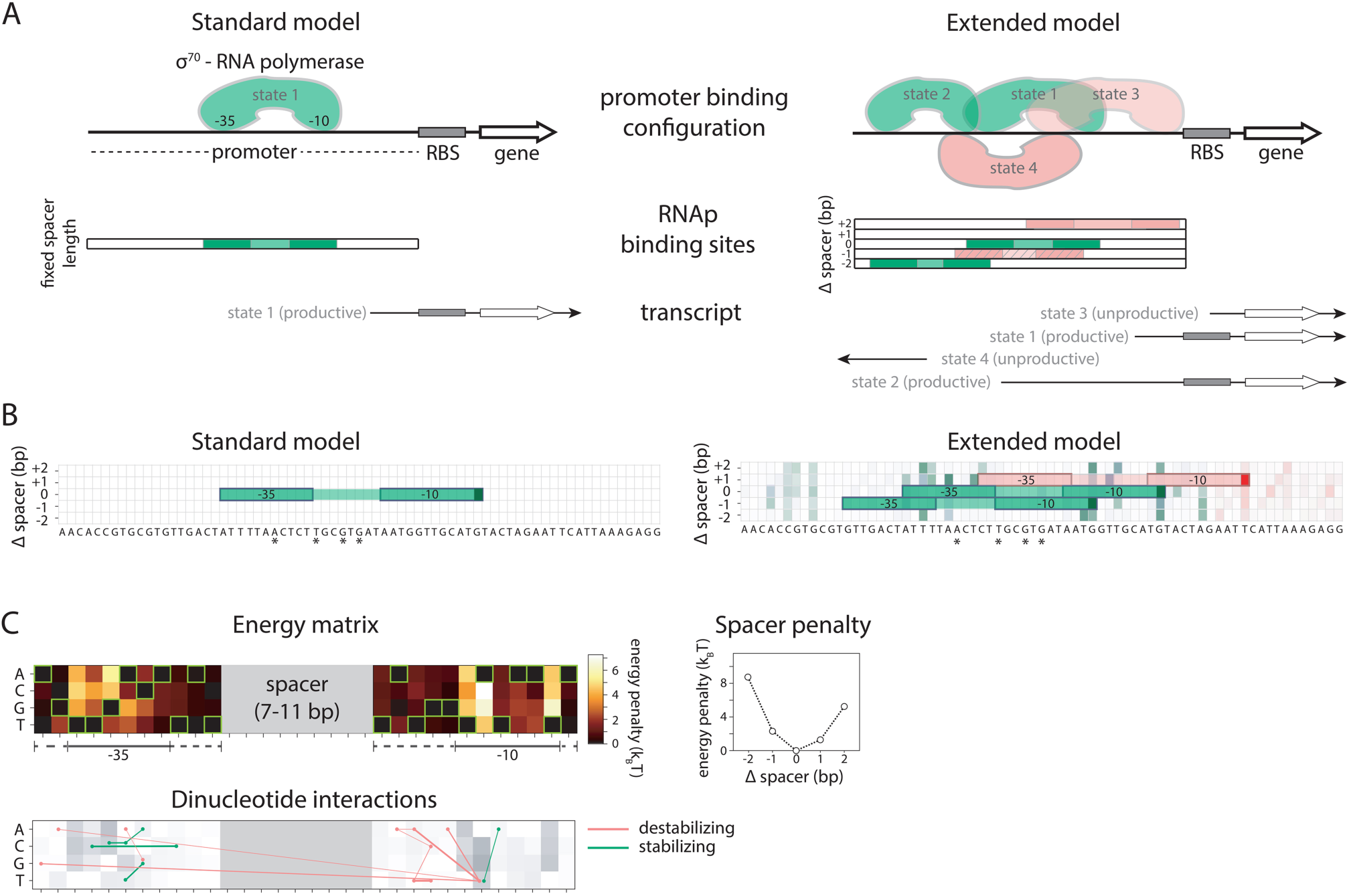
(**A**) The standard thermodynamic model assumes only one (strongest) σ^70^-RNAP binding configuration at the promoter, which generates a productive transcript. We refer to ‘promoter’ as the entire *cis*-regulatory element, while ‘binding site’ refers to the RNAP contact residues of a specific binding configuration (colored area in ‘RNAP binding sites’). The Extended model incorporates structural features of bacterial promoters into the thermodynamic framework: (i) the possibility that σ^70^-RNAP binds a single promoter region in multiple configurations that independently and cumulatively contribute to expression (*25, 26*), as observed in transcriptomics studies (*27*) (binding states 1-4); (ii) spacer length flexibility, captured by an energy penalty for changing the distance between -10 and -35 elements away from the optimal (difference between states 1 and 2); (iii) occlusive unproductive binding, which occurs when the initiation start site excludes a part of the ribosomal binding site (RBS) resulting in a non-translated transcript (state 3); (iv) occlusive binding on the reverse complement, which inhibits productive binding at the promoter (*28*) (state 4); (v) dinucleotide interactions between promoter residues that are in direct contact with σ^70^-RNAP; and (vi) clearance rate of RNAP from the promoter, which captures, in a very simplified manner, the complex events that occur on the DNA before σ^70^-RNAP is released from its binding site (v and vi not illustrated in figure). We experimentally verified features (i), (iii) and (iv), as they have not been characterized before (Fig.S1-3). We do not account for the UP element (*21, 29*). (**B**) Example model output for a selected *P*_*R*_ mutant (stars mark mutated positions) for the Standard and the Extended model, showing binding configuration on the forward strand. Pixels forming the background grid indicate -10 end-points of binding sites, with the intensity of color corresponding to the strength of binding in that configuration (green productive, red unproductive binding). States that are bound strongly enough to independently lead to measurable expression are framed (one for the Standard model, three for the Extended model – two productive, one unproductive). (**C**) Main biophysical parameters fitted from *sort-seq* data. Energy matrix shows the effect of every possible binding site residue on the binding energy between σ^70^-RNAP and DNA (strongest binding indicated by green squares). The optimal energy matrix consists of the -10 and -35 elements (underlined), positions outside the canonical elements that significantly affect quantitative predictions of gene expression levels (dotted underline), and spacer of optimal length 9bp (corresponding to the canonical 17bp between -10 and -35 elements). Strongest stabilizing (green) and destabilizing (red) interactions between dinucleotides are shown, with line thickness indicating the deviation from independent energy contribution to binding (range 0.15-0.38k_B_T). For other model parameters and all significant dinucleotide interactions, see Table S1,S2.

Both, the Standard and the Extended model, were fitted by subsampling a library of >12,000 constitutively expressed random mutants of one of the strongest known promoters, Lambda *P*_*R*_, that controlled the expression of a *yfp* reporter gene from a small copy number plasmid in *E.coli* (Fig.S4A). *Sort-seq* experiments were used to measure the gene expression level of mutants (*11*): mutants were separated using a cell sorter into four phenotypic bins (‘no’, ‘low’, ‘medium’ and ‘high’ expression), PCR-tagged according to the sorted bin, and bulk sequenced to obtain the counts of each mutant sequence in the sorted bins (Fig.S5). To infer model parameters, we calculated the binding strengths of σ^70^-RNAP to DNA in all possible configurations allowed by the Extended model. Then, we integrated over all positive (‘on’) and negative (‘off’) configurations to obtain the probability (P_on_) of productive σ^70^-RNAP binding, which solely determines expression levels. We searched for optimal model parameters (Fig.1C) by maximizing the likelihood of the weighted multinomial logistic regression of log_10_P_on_ to the median observed expression bin for a training subset (40%) of the *P*_*R*_ mutant library. The model reproduced the evaluation subset of the *P*_*R*_ mutant library with high accuracy, significantly improving on the performance of the Standard model (Fig.2B).

**Figure 2.**
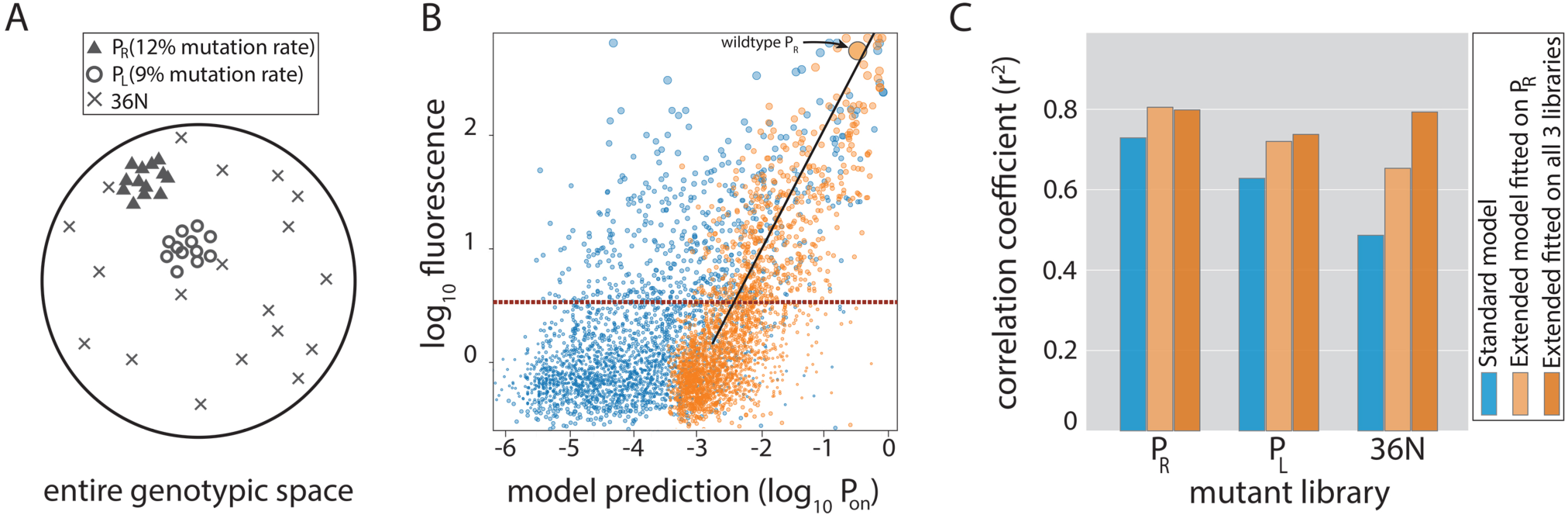
(**A**) Cartoon of the three mutant libraries: *P*_*R*_ and *P*_*L*_ sample locally around a wildtype; *36N* randomly samples the full 36bp-long genotypic space. (**B**) Model-to-data correlation for the Standard (blue) and the Extended model (orange) trained on *P*_*R*_ only, shown for the evaluation subset (20%) of the random (*36N*) mutant library. Line of best fit (solid black) and the instrument detectability threshold (red dashed line) are shown. Marker sizes indicate data-point weights used in fits. (**C**) Model performance on mutant libraries (*P*_*R*_, *P*_*L*_, *36N*), shown as fraction of variance explained on evaluation data.

To assess the cross-dataset predictability of the two models, we created two additional mutant libraries and analyzed them in a *sort-seq* experiment (Fig.2A). One of the libraries consisted of random mutants of the Lambda *P*_*L*_ promoter, which shares sequence homology to *P*_*R*_. To determine model performance across the entire unconstrained genotypic landscape, we built a third library consisting of completely random 36 nucleotides controlling the expression of a *gfp* reporter gene (‘*36N*’). This library was created in a different strain (Fig.S4B), and the expression was sorted into 12 bins instead of 4, giving greater precision in testing the model-to-data fit. Interestingly, ∼10% of experimentally-measured, random 36bp-long sequences led to measurable expression (Fig.S6).

While the Extended model modestly improved predictability of the *P*_*R*_ and local cross-predictability (*P*_*L*_) datasets, it dramatically improved predictions of gene expression levels from random sequences (Fig.2B,C). Each of the structural features included in the Extended model (Fig.1A) significantly improved predictability (Table S3). Fitting the Extended model by subsampling all three libraries (>25,000 mutants) led to further improvements in predictability of expression from random sequences (Fig.2C), while also allowing more reliable estimation of dinucleotide interactions between promoter positions that contact σ^70^-RNAP (Fig.1C). Dinucleotide interactions are moderate contributors to promoter function (Table S3) (*12*), and appear to be stronger when at least one residue is outside the canonical -10 and -35 binding sites.

Low cross-predictability of the Standard thermodynamic model (Fig.2C) is often attributed to context-dependence: the poorly understood effect of positions flanking the -10 and -35 elements (*8, 9*). The Extended model explains away the extensive context-dependence as a consequence of the promoter structural features that we account for (Fig.1A).

Having a biophysical model that accurately predicts gene expression levels from random sequences enabled us to sample the entire genotypic space in order to describe the genotype-phenotype mapping of constitutive promoters. Approximately 20% of random 115bp-long sequences (average inter-genic region length in *E.coli*) are predicted to have measurable expression (compared to ∼8% for the Standard model) (Fig.3A). Surprisingly, for ∼82% of non-expressing random sequences, at least one point mutation could be found that led to measurable expression (Fig.3B) – a finding that we verified experimentally (Fig.S7). In total, more than 1.5% of all possible point mutations did so (Fig.3C) indicating that promoter sequences readily emerge, as previously suggested based on a limited sample of 40 sequences (*13*).

**Figure 3.**
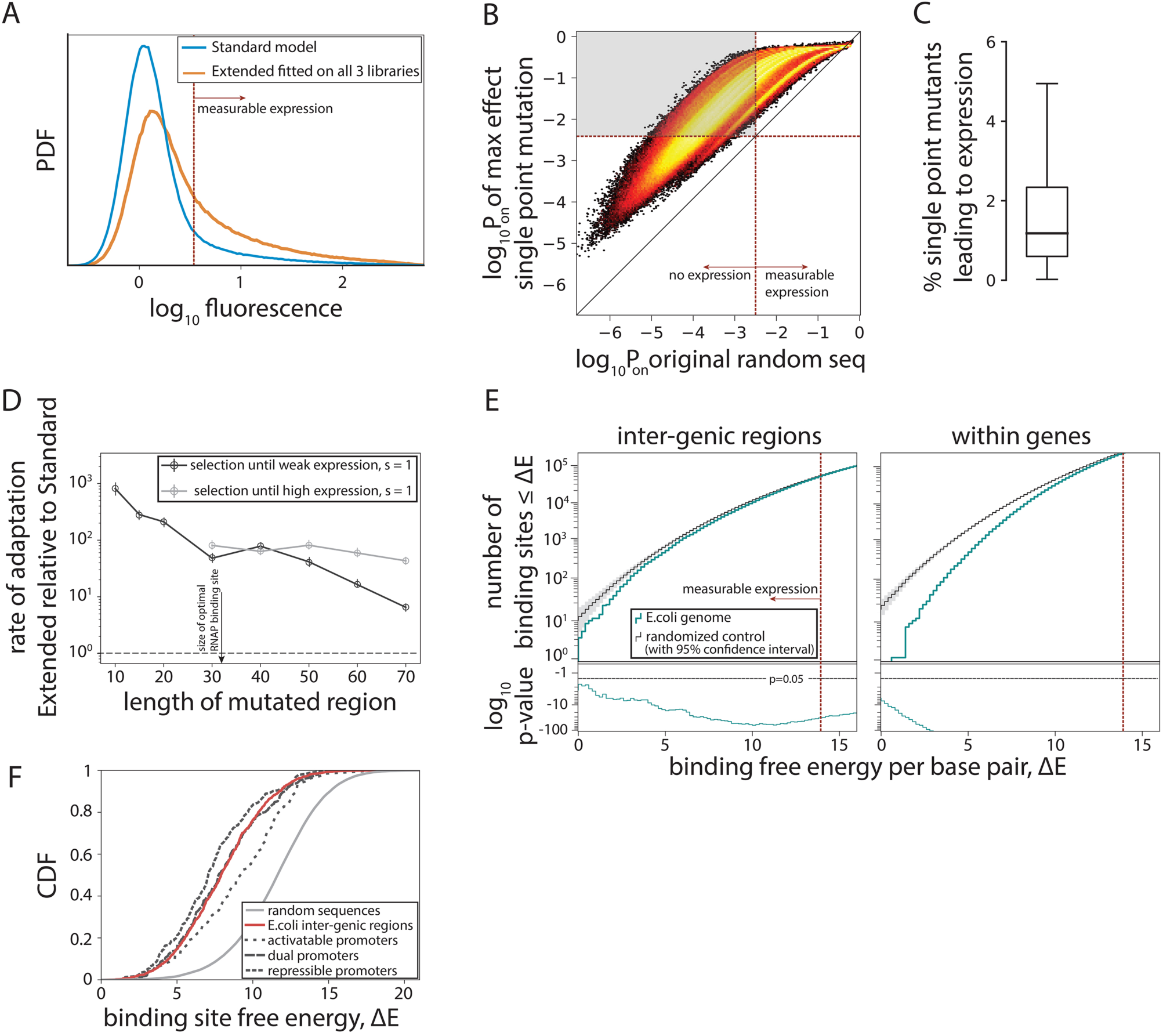
(**A**) Probability density function (PDF) of simulated flow cytometry fluorescence intensities from 10^6^ randomly generated 115bp-long sequences. Red dotted line marks the cutoff for ‘measurable expression’, fitted from experimental data. (**B**) Density heat map (brighter color represents higher density), showing, for every simulated random sequence (expression on x-axis), the expression of a single point mutant with the largest positive effect on predicted expression (P_on_) (y-axis). For 82% of non-expressing random sequences (sequences left from the dotted line on the x-axis), that mutation led to measurable expression (gray area). (**C**) Box-plot showing the percentage of all possible point mutations that convert a given random non-expressing sequence into one with measurable expression (obtained from 10^5^ random sequences). (**D**) Increase in rates of evolution of the Extended relative to the Standard model. Evolution to either weakest measurable expression, or high (*P*_*R*_) expression levels was modeled through single point mutations. Evolution was simulated 100 independent times for each of the 100 random 115bp-long starting sequences by mutating the central contiguous part of the indicated length. Evolving promoters would almost never reach high expression levels when only a region smaller than the RNAP binding site (30bp) was allowed to mutate. (**E**) For evidence of selection against σ^70^-RNAP binding sites we compared the free energy per bp between either the inter-genic (promoter containing) or the within-genic regions of the *E.coli* genome and that of a random sequence with the GC% of the corresponding region (note that higher energy means weaker binding and hence lower expression). Associated p-values are also shown. (**F**) Cumulative distribution functions (CDFs) for predicted binding strengths of different *E.coli* promoters, obtained from RegulonDB.

To contextualize these findings, we simulated evolution under directional selection for expression from random non-expressing sequences using a Strong-Selection-Weak-Mutation model (*14*) adapted from Tugrul *et al*. (*15*). The Extended model predicts rapid promoter evolution, establishing the upper speed limit for the emergence of new functional promoters (Fig.S8). Constitutive promoters evolve rapidly from non-expressing sequences because of the structural promoter features (Fig.1A), which dramatically enlarge the number of possible binding configurations that contribute to expression – a phenomenon particularly evident at lower expression levels, where cumulative binding across multiple binding sites substantially contributed to overall expression (Fig.S9). For these reasons, the Extended model predicts, on average, more rapid evolution from a non-expressing sequence towards both, weakest measurable expression and strong (wildtype *P*_*R*_ level) expression when compared to the Standard model (Fig.3D, S10). The difference in rates of evolution becomes even more pronounced as the mutated sequence gets shorter (simulating stronger constraints due to, for example, existence of other transcription factor binding sites) (Fig.3D).

The pervasiveness of σ^70^-RNAP binding sites among random DNA sequences has surprising implications for understanding the evolution of gene regulation: the critical question is not how promoters arise, but rather how a cell avoids expressing everything, all the time. To address this, we asked if there was evidence of selection against σ^70^-RNAP binding sites in the *E.coli* genome (Fig.3E, S11) (*16*). While identifying strong selection against σ^70^-RNAP binding sites within genes (*13*), surprisingly, we also found evidence of selection against binding sites in inter-genic regions (p<10^−40^ across all inter-genic regions). Thus, there is selection against σ^70^-RNAP binding sites in the *E.coli* promoter regions themselves. This negative selection was observed even when only the known σ^70^ promoters (obtained from RegulonDB (*17*)) were considered (Fig.S12), most of which strongly bind σ^70^-RNAP and hence lead to intermediate and high expression levels (Fig.3F).

Mapping promoter genotype to gene expression phenotype is fundamental for understanding gene regulatory network structure and evolution (*18, 19*) and engineering synthetic biological systems (*20*). Accounting for structural features of promoters enabled us to accurately predict gene expression levels of any DNA sequence, extending promoter genotype-phenotype mapping beyond local neighborhoods of known promoters (*8, 9*) to capture the entire genotypic space. Recently, a complementary approach described two further structural features of DNA-RNAP interaction, the avidity between -10 and -35 binding sites and the UP element (*21*), yet was not designed to predict expression from arbitrary sequences. Integrating these features into our Extended model could further improve promoter sequence-to-function predictions. Furthermore, the thermodynamic framework can be extended to model complex, regulated promoters (*7, 22*).

The flexibility of σ^70^-RNAP binding ensures the proximity of any sequence to a constitutive promoter (Fig.3). This allows σ^70^-RNAP to function as a global regulator (*17, 23*), permitting selection to sustain binding specificity of required transcription factors and reduce crosstalk while maintaining expression (*24*). The ease of evolving constitutive promoters, which we observed using large random mutant libraries, has been experimentally hinted at (*13*) but remained theoretically unexplained (*15*). Developing biophysical models based on molecular properties of biological systems can reconcile experiments with theory, and form a critical framework for understanding how organisms function and evolve.

## Supporting information

all supplementary figures and tables

## Acknowledgments

We thank Hande Acar, Nicholas H. Barton, Rok Grah, Tiago Paixao, Maros Pleska, Anna Staron and Murat Tugrul for insightful comments and input on the manuscript.

## Funding

This work was supported by: Sir Henry Dale Fellowship jointly funded by the Wellcome Trust and the Royal Society (Grant Number 216779/Z/19/Z) to M.L.; IPC Grant from IST Austria to M.L. and S.S.; ERC Advanced Grant (250152); and European Research Council Funding Programme 7 (2007-2013, grant agreement number 648440) to J.P.B.

## Author contributions

M.L., S.S., M.S., G.T, C.C.G. conceived the study. M.L., M.S., and D.T-A. performed the experiments. S.S., G.T., and M.L. analyzed data. S.S. and G.T. performed the formal analysis and developed the model. M.L. wrote the original manuscript and revised with S.S., M.S., D.T-A., J.P.B., G.T. and C.C.G.

## Competing interests

Authors declare no competing interests.

## Data and materials availability

All data and code will be made available upon publication on a free-to-access public repository (Dryad and GitHub), and is available upon request from authors.

## Materials and Methods

### Experimental systems and mutant libraries

The *P*_*R*_ system consisted of *venus-yfp* (*30*) under the control of Lambda bacteriophage *P*_*R*_ promoter (Fig.S4A). The system was isolated from the rest of the plasmid with two terminators, T1 and T17. The *O*_*R3*_ site of *P*_*R*_ was removed in order to remove the *P*_*RM*_ promoter. The ribosomal binding site (RBS, carrying the Shine-Delgarno AGGAG sequence) was 28bp away from the downstream end of the strongest σ^70^-RNAP binding site in the *P*_*R*_ promoter. The entire cassette was inserted into a low-copy number plasmid backbone SC101* carrying a kanamycin resistance gene (*31*). The random *P*_*R*_ mutant library was created by cloning custom-made oligonucleotides (IDT Technologies), which had a 12% mutation rate for each of 67 positions in the *P*_*R*_ promoter (4% mutation chance for each possible mutation away from the wildtype), instead of the wildtype promoter. Ligated plasmids were electroporated into One-Shot^®^ Top10 cells (Life Technologies, Carlsbad, US). This step was used to maximize the library diversity due to One-Shot^®^ Top10 cells’ high competency. Following electroporation, cells were grown for 1h in LB and then plated on selective kanamycin plates to allow single colony formation and minimize resource competition, and grown overnight. To ensure large coverage, we cloned mutagenized PCR products until we obtained at least 30,000 individual colonies (uniquely transformed individuals). Using chilled LB media, colonies were washed off plates and collected. Plasmids were isolated from this collection in bulk, and cloned into strain MG1655-K12. The same wildtype layout and mutagenesis protocol was used to create the *P*_*L*_ mutant library, with the exception that mutation rates were 9% per nucleotide (with 3% mutation chance for every possible non-wildtype mutation). Cells were always grown at 37°C.

The *36N* library was placed in pUA66-lacZ plasmid backbone carrying Kn resistance (*32*), by cloning a 100bp-long oligonucleotide containing 36 random base pairs (each of the 36 positions had a 25% chance of being either adenine, thymine, cytosine or guanine), surrounded on each side by 32bp of randomly-generated, non-expressing DNA sequence (Fig.S4B). The two 32bp-long segments had a different sequence. This 100bp sequence was placed upstream of the RBS, controlling the expression of a *gfp* gene (*32*). The *36N* plasmid DNA library was generated using a Q5 site directed mutagenesis kit (New England Biolabs, Ipswitch, MA). For amplification, we used the reference plasmid as a template (a plasmid with a random, non-expressing 100bp fragment) and two pools of primers with a constant 3’ end and an 18N random 5’ end. We cloned the mutagenesis products into electrocompetent NEB5α cells. Following electroporation, cells were grown for 1h in LB and then plated overnight on selective kanamycin plates. Cells were collected in bulk to form the *36N* mutant library. We used different strains in order to better evaluate the generalizability of our model. The mutation rates of all three libraries (Fig.S13), as well as the sequence-level randomness of the *36N* library, were checked based on the *sort-seq* data.

### Sort-seq experiments

Prior to sorting, cells were grown in M9 minimal medium with 0.2% CAS, 0.2% glucose and 50 μg/mL kanamycin. Frozen aliquots of the mutant library *36N* library were diluted 1:10 and grown overnight. Prior to sorting, overnight cultures were diluted again 1:100 and grown for 3h to reach exponential phase. We repeated the sorting for 3 biological replicates.

FACS sorting was performed on an FACS Aria III flow cytometer (BD Biosciences, San Jose, CA) with a 70μm nozzle for droplet formation. A 488nm laser was used to detect forward scatter (FSC) and side scatter (SSC) with a 488/10 band-pass filter. FITC channel was used for excitation of either YFP (*P*_*R*_ and *P*_*L*_ libraries) or GFP (*36N* library). The flow rate was set to 1.0 and samples were diluted to obtain a cell count of approximately 2000 events/second. Cells for sorting were manually gated on the densest population in an FSC/SSC scatter plot, which comprised 95.5% of all events exceeding a threshold of 1000 on the SSC axis. For the *P*_*R*_ and *P*_*L*_ libraries, 4 sorting gates were set on FITC: no-expression gate, capturing >99% of all measurements from a non-expressing plasmid (control plasmid not containing *yfp*); high-expression gate, capturing >99% of all measurements from the wildtype *P*_*R*_ plasmid; two gates equidistant in fluorescence between the no- and high-expression gates (Fig.S6). The *36N* library was sorted into twelve gates as follows: The upper boundary of the lowest gate corresponded to the median of an auto-fluorescence control sample (plasmid-free cells). The lower boundary of the highest gate (B12) was set to 2×10^4^. Distances between the remaining intermediate nine gate boundaries were of equal size on the log-scale FITC histogram (Fig.S6). For *P*_*R*_ and *P*_*L*_ libraries we could sort into all four bins simultaneously, and hence we sorted 1 million cells for each of the three biological replicates. For the *36N* library, we first recorded 10^5^ reads, and then the number of cells sorted into each of the twelve bins corresponded to the number of cells recorded in each of the bins. The recipient plate was cooled to 4°C to halt growth while sorting to other wells was still going on. Only for the *36N* library, after sorting we added 1000 cells with the reference plasmid into each of the 12 sorted populations. We did this to maximize the precision of our experimental measurements for the *36N* library, as it would enable more accurate experimental determination of expression levels by enabling normalizing the number of mutants in each bin (see section *Processing of the 36N mutant library*).

Cells from each sorted bin were grown overnight. We isolated plasmids from the sorted populations. We used high-fidelity PCR (Phusion, New England Biolabs) to amplify 150bp containing the mutagenized region, and barcoded the primers according to the sorted bin. Four sets of barcoded primers were used for the *P*_*R*_ and *P*_*L*_ libraries, and 12 for the *36N* library. PCR products were column-purified (Zymo Research, Irvine, CA) and eluted in 30 μL, of which 2 μL were run on an agarose gel for product quantification based on band fluorescence. PCR products were pooled to reach approximately equimolar concentrations of each bin, separately for each mutant library. No additional clonal amplification steps were conducted prior to sequencing. Each library was sequenced with millions of reads using 135-bp pair end Illumina sequencer (Hi-seq).

### Processing of P_R_ and P_L_ mutant libraries

Here, we describe the data processing pipeline for the *P*_*R*_ and *P*_*L*_ libraries, from the initial sequence reads to a dataset we use for fitting and evaluating out model. For each library, we obtained millions of reads (Table S4), which we paired (using illumina-utils package (*33*)), discarding reads with any mismatch. Each read contained a tag on both ends with information of the expression bin the sequence was sorted in in the FACS, and we only took reads which have the same tag on both ends.

In each library, the remaining ∼2.5 million reads contained more than 300,000 unique sequences, which we filtered based on length, coverage, and the position of the ribosomal binding site (Shine-Dalgarno sequence that was the same for all sequences in the library as it was not mutated) (Fig.S14A,B). We further required the sequences not to be too different from the ancestral sequence (*P*_*R*_ and *P*_*L*_ respectively) (Fig.S14C). This step removed the remaining *P*_*R*_ sequence and a small cloud of sequencing errors around it from the *P*_*L*_ library. *P*_*R*_ wildtype sequences were present in the *P*_*L*_ library because both libraries were made using the *P*_*R*_ wildtype sequence as the starting point for cloning and ligation – a technique that is nearly but not exactly 100% efficient. Even though we used high-fidelity polymerase for the PCR, still some sequencing errors existed (as evident from the ‘cloud’ of sequencing errors around the *P*_*R*_ wildtype in the *P*_*L*_ library), but were lower than 0.01%. Finally, we required the distribution of expression bins to be as unambiguous as possible (Fig.S14D,E).

We observed a discrepancy in the number of unique sequences in the two libraries (Table S4) that arose due to the higher mutation rate of the *P*_*R*_ library. This is why we took a more conservative approach and additionally raised the coverage threshold to 30 for all our analyses.

### Processing of the 36N mutant library

The original sequences library contained more than 10 million paired reads, almost all of which paired without mismatches (Table S5). Out of those, more than a million contained the reference plasmid sequence, which we extract and treat separately. We required sufficient mapping (higher than 0.75 similarity, using local alignment function from the pairwise2 module in Biopython, with symmetric gap open and extension penalty of − (2L)^−1^, and a matching score of L^−1^, where L is the length of the mapping region) of both flanking regions, and that the length of the core region is within 2bp of the canonical (36bp). Most of those reads were unique and most probably errors, so we originally required coverage of at least two (Table S5).

At this point we set to estimate the abundance of spurious sequences (e.g. sequencing or PCR errors), by cross-mapping the abundant sequences (with the highest coverage) against all other sequences, using the same scores as defined above (Fig.S15A). In every case, the distribution of mapping scores is bimodal, with an overwhelming number of low scores and a small number of very high scores, suggesting that abundant sequences come with a cloud of errors, which are necessarily similar to the investigated sequence. Compiling a distribution of similarity scores for the 1000 most covered sequences, showed a clear threshold of 0.7 between the low- and high-scored modes. We then accumulated all the sequences that appeared to be associated with a number of high-covered sequences, and noted that the histogram of coverages of only those sequences closely matches the low-end part of the histogram of all sequences. Extrapolating from this, we estimated that requiring a coverage of at least 30 reads would lower the number of spurious sequences to only a few dozen (Fig.S15B).

In contrast to the *P*_*R*_ and *P*_*L*_ libraries, where FACS sorting was performed simultaneously into four bins, here the experiment involved sequential sorting of the same number of cells into one of the 12 bins (Fig.S6). Naturally, less abundant expression bins had to be sorted for a longer time, which introduced a systematic bias for higher expressing bins. To account for this bias towards higher-expressing sequences, we introduced a known number of a reference sequence to each bin.

To debias, we divided each distribution by the distribution for the reference sequence (Fig.S15C), and normalized. We then constructed a set of average distributions for all sequences that have the same mode. This allowed us to fit the background noise (intrinsic to FACS measurements) for all distributions. Next, we cleaned the sequence-specific distribution of potential outliers that may distort the estimate of the mean: for each mode-specific template distribution, we defined a filter that selected only bins in which the background is at max ⅓ of the value of the template. For each sequence-specific distribution, we selected a filter based on its mode, which nullified values in bins defined as outliers (Fig.S15D,E). Afterwards, we renormalized the distribution. This filtering is especially important for the higher-expressing sequences, where a single count in the lower bins would get enlarged tremendously by debiasing, drastically skewing the expression estimate (Table S5).

We used the debiased and filtered distribution over bins *α*_*i*_ = [*α*_0_, …, *α*_11_], to produce two estimates of expression. In the units of bin index, we estimate expression as; 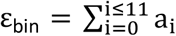 while in the units of luminosity measured in FACS as 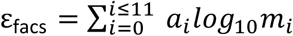, where m_i_ is the median value of measurements in the i-th bin. The two estimates are linearly related.

To validate out data processing, we randomly picked 8 mutants from each of the 12 bins, sequenced them and measured in a plate reader. Out of 96, 79 were unique mutants, and two did not exist in the *36N sort-seq* library. We compared the measurements of the remaining 77 sequences obtained from a plate reader to the estimate of their expression obtained from the *sort-seq* experiment following the above-described debiasing and filtering (Fig.S16). Plate reader measurements were conducted in the following manner: Mutants were grown overnight in M9 minimal media supplemented with 0.2% CAS, 0.2% glucose, and 50μg/mL kanamycin. The overnight populations were diluted 1,000-fold, grown until OD_600_ of approximately 0.1, and their fluorescence measured in a Bio-Tek Synergy H1 platereader. Three replicates of each mutant were measured.

### Data splits and fitting procedure

We define 60:20:20% splits of each of our libraries into three disjunct datasets (Table S6, S7, S8, S9). The first (‘Training’ dataset) and second (‘Validation’) were used for training and model selection respectively, while the last (‘Evaluation’) was used exclusively for final evaluation and visualization of our models.

The central quantity of our modelling approach is the proportion of time RNAP spends bound to the DNA in the “on” configuration, P_on_ i.e. in a configuration that can yield productive mRNA and further leading to a protein. Our quantification of expression, which we define as log fluorescence, will then be proportional to log_10_ *P*_on_ (*7, 10*).

The three libraries were different in terms of their output. While for the *P*_*R*_ and *P*_*L*_ libraries we used only four bins and chose the median bin for each mutant (more conservative statistic), we obtained more plausible estimates of expression for the mutants in the *36N* library (as explained above). To keep the procedure the same across all three libraries, for the *36N* library we rounded the estimate of the mean expression in bin units to the closest integer. This allowed us to use multinomial logistic regression for all three datasets and the associated log-likelihood as the objective function.

Concretely, given a set of parameters (binding matrix, spatial penalties, etc.), our model produces log_10_ *P*_on_ for each sequence, which we use as an independent variable to fit observed bin expression levels using multinomial logistic regression (from scikit learn, with L-BFGS-B optimization (*34*)). Additionally, we explicitly required a balanced fit by applying weights inversely proportional to the number of observations in each bin for each dataset. This is especially important for the *36N* library, due to the highly disproportionate numbers of observations in the 12 bins.

The main metric for our optimization and model performance is the likelihood of the logistic regression. In the interest of higher interpretability, we also report the *r*^2^ value of a linear fit where the independent variable is log_10_ *P*_on_ and the dependent variable is the log fluorescence estimate (median for the *P*_*R*_ and *P*_*L*_ libraries, as a robust measure in the absence of high bin resolution; and mean for the *36N* library, as having 12 bins allowed us to accurately estimate the mean), using the same weights as for the logistic regression. The multinomial logistic regression does not necessarily yield a linear dependence between log_10_ *P*_on_ and observations. In fact, across the range of reasonable values, the log-likelihood depends on chemical potential only weakly, yet it may change the correlation coefficient by several percent. For this reason, in Fig.2C, we show correlation coefficients (*r*^2^) for evaluation datasets with chemical potentials reoptimized on the respective training datasets. Raw *r*^2^ values are similar and are reported in Table S3.

We start from the Standard model, and progress towards the Extended model sequentially, including one structural feature at the time. We fit the model parameters using only the training dataset of the *P*_*R*_ library, and only searched in the vicinity of the previous best fit. We assess such a model on the *P*_*R*_ evaluation and all three datasets of the *P*_*L*_ library without any adjustments, since the two libraries were obtained following the same experimental protocol in the same cells. In that sense, the whole *P*_*L*_ library played the role of an evaluation set. To evaluate this model on the *36N* library, which was obtained using a different cell line and through different FACS thresholds, we used the *36N* training set to re-fit the chemical potential and the hyper parameters of the logistic repression. Therefore, in the context of evaluating a model fitted on the *P*_*R*_ library alone, both validation and evaluation *36N* datasets can be considered true validation datasets.

Ultimately, we wanted to pool the training datasets of all three libraries to fit a unique model to give us the best set of parameters. In this case, all validation and evaluation datasets were data-naive. Doing this increased the inference power of our models, allowing us to fit pairwise (dinucleotide) interactions between nucleotides and the rate of RNAP clearance from the initial binding site.

### Standard thermodynamic model

In thermodynamic models of gene expression, the amount of protein is directly proportional to the fraction of time RNAP is bound to the promoter sequence. In the simplest case, where RNAP can be either bound (*“on”*) or non-bound (*“off”*-state), the probability of the *on*-state is given by the formula:

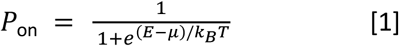

where *E* is the binding energy, and *μ* the chemical potential of the RNAP in the cytoplasm. All our experiments were performed at room temperature, so *k*_*B*_T = 0.59 kcal/mol. For simplicity, we define energy and chemical potential in those units, and drop *k*_*B*_T from subsequent formulas.

We further expressed binding energy as an independent sum of local interactions between RNAP and individual base pairs in the promoter region, with each of the four possible bases (A,C,G,T) contributing differently depending on the position in the promoter. Such 4 × *l* matrix is referred to as the binding Energy Matrix. The values of the Energy Matrix are determined up to an arbitrary offset per position. We set zero at the binding energy of the wildtype *P*_*R*_ sequence.

In the case when the promoter sequence L is longer than the binding matrix l, the number of states is N = L − l + 1, and the *on*-probability takes the form

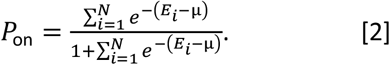

The Standard model assumes that only one element in the sum dominates, so we follow other studies (*7, 10*) when formulating our standard thermodynamic model as

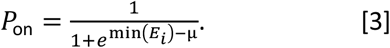

### Towards the Extended model: spacer flexibility

Based on the common observation that RNAP has a flexible spacer length, we extended the Standard model by allowing for spacer lengths that differ from the canonical by up to 2bp. This effectively increased the number of possible configurations by a factor of five. The key difference that we introduced was to assign an energy penalty for each non-canonical configuration. Increasing the spacer flexibility beyond 2bp did not yield further benefits to predictability.

### Towards the Extended model: cumulative binding

In order to account for multiple RNAP binding configurations (different positions along the promoter and different spacer lengths) that can lead to a productive transcript, we saw the necessity of actually performing the thermodynamic sum in Eq.[2] instead of just extracting the dominant binding as was done for the Standard model. Fully embracing the thermodynamic description at this point provided us a natural language for all further extensions.

### Towards the Extended model: occlusive unproductive binding

We accommodated the occlusive unproductive binding states naturally in the thermodynamic description:

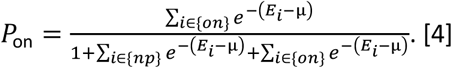

The main challenge here was to find the exact position after which the transcripts were produced without the RBS. To answer this question, we checked computationally whether there was a natural region where this would be the case. First, we aligned all sequences with respect to the position of the ribosomal binding site (Shine-Dalgarno sequence “AGGAG”). Then, using the parameters from the previous model iteration (with cumulative binding), we assessed performance with different positions separating the productive and unproductive binding. This identified an 8bp-long region, where the separating position was to be expected (Fig.S17). To determine the location more precisely, we set up a separate experiment (see section *Verifying promoter structural features*).

### Towards the Extended model: occlusive binding on the reverse complement

When bound to the reverse complement, we considered RNAP to effectively act as its own repressor. We accommodated this in the model by expanding the set of “np” states in Eq.[4] to all reverse-complement configurations. Including this effect led to a slight but significant increased performance even without refitting, after which we also re-optimized the parameters locally.

### Towards the Extended model: dinucleotide interactions

The previous components of the model were fit exclusively on the *P*_*R*_ dataset. In our search for dinucleotide interactions and when exploring departures from thermodynamic equilibrium, we pool the datasets in search for the best fit, as only then could we get a significant estimate of the interactions.

A common assumption of thermodynamic modelling of gene expression is that the binding energy of any particular transcription factor-promoter complex can be expressed as an independent sum of interactions between individual base pairs and the transcription factor (RNAP in our case) residues. A naive inclusion of all possible dinucleotide interactions between the contact residues of RNAP and the promoter would inflate the number of parameters to thousands, rendering their simultaneous estimation hopeless, even before considering overfitting issues.

To overcome the pitfalls of overfitting and estimate the potential importance of dinucleotide interactions, we first included each interaction independently, and required that the best fit increased log-likelihood by at least 2. This reduced the number of interactions from 4,416 to 892. Then, we drew 10 random subsets of 20 interactions, and jointly optimized the interactions in the vicinity of the previously obtained values, over 10 cross-validation splits (50:50) of the training and validation datasets and using L1 regularization. This way, we obtained estimates (and associated error) for the interaction value in 10 different “backgrounds”. For each interaction, we considered valid only those estimates that were larger in magnitude than a defined threshold, and combined them into a single estimate (mean and standard deviation), allowing for a small amount of leniency due to rather strict L1 regularization. We chose 0.002k_B_T as the threshold for acceptance and made sure the results were robust for thresholds of 0.001 and 0.005. We then filtered for only those interactions that were non-compatible with 0 at confidence level of 2*σ*. This brought down the number of interactions to 250. In the next step, we again jointly inferred subsets of interactions: this time 12 subsets of 50 interactions. For each interaction, we then estimated the single mean and standard deviation from the 12 values, and conservatively selected only those that were inconsistent with zero at more than 3*σ*, which brought down their number to 77. Finally, we sequentially include dinucleotide interactions starting with the most and moving towards the least significant, requiring that each contributed to at least 3*σ* -level in the background of all accepted up to that point. This left 31 interactions (Table S2).

### Towards the Extended model: clearance rate

Finally, we push our framework even closer to reality by noting that RNAP is not like any other DNA binding molecule, in-so-far that it is in fact *not* in thermodynamic equilibrium: to make the transcript, it needs to leave its original binding site. We model this by introducing a parameter R, as a rate with which RNAP is cleared away from the binding site, relative to the rate with which it is bound, in the limit of very strong binding. It effectively sets up an upper limit on the amount of time RNAP persists on the binding site, and should make the difference only with strong binding sites.

Consider a system where RNAP can exist in only three states: 1) bound at a productive position, 2) bound at an unproductive position, and 3) unbound (“off”); and consider the

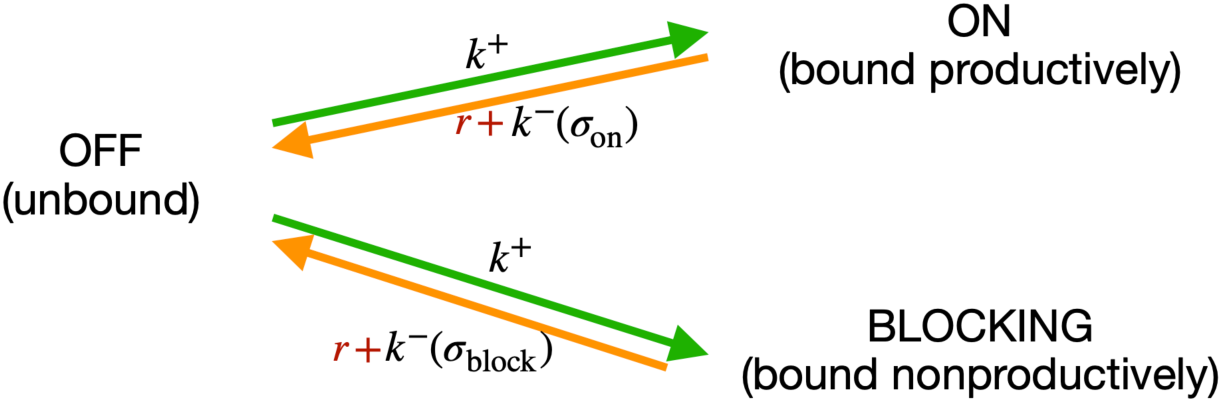

transition rates among them:

The transition rate from unbound to any of the bound states *k*^+^ is the same and depends only on the concentration of free RNAP in the vicinity of the sites. The reverse transitions depend on the sequence (σ), through energy of binding 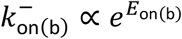. We modeled RNAP clearance as an independent rate *r*, which depletes the bound states *on* and *b*, and eventually contributes to repopulation of RNAP in the cell. We do not model time dependence, and are interested only in the stationary state, thus we do not need to model the time delay between leaving the bound state and reappearing in the cell.

In stationarity we have

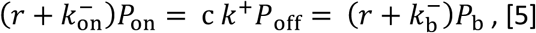

Introducing a relative clearance rate 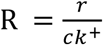, casting the known rates and RNAP concentrations in terms of binding energy 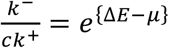, and considering that *P*_on_ + *P*_b_ + *P*_off_ = 1, we obtained:

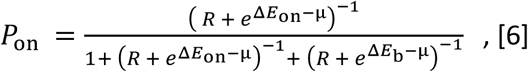

This is the same formula as Eq.[4], with 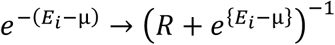. Considering clearance rate effectively introduced a cutoff on the binding energy *E*_min_ = *μ* + log R, below which the energy of binding is not important anymore. Note that in the case of R → 0, the formula reduces to the previous state. In the case of a single (on) binding site, *R* is a completely degenerate degree of freedom, which can be manifestly removed by a transformation *μ* → *μ* − log(1 + *R*).

The scan over the values of *R* (only on training dataset) is shown in Figure S18. For each value, we refit the chemical potentials. We obtained best fit value for the relative clearance rate of *R\** = 0.31, and a large uncertainty region, yet the model quite robustly preferred a non-zero clearance rate. Here, we consider clearance rate a property of the RNAP molecule itself, and independent of sequence. Including the sequence dependence for the clearance rate can be done, but disentangling the exact nature of such dependence would require a dedicated set of experiments that is beyond the scope of this work.

### Verifying promoter structural features

We incorporated several structural features of bacterial promoters into the thermodynamic modeling framework (Fig.1B). When estimated on all three mutant libraries, each of these features independently increased model predictability (Fig.2C), providing evidence of its biological role. Nevertheless, we conducted additional, hypothesis-driven experiments to verify the role of these structural features. For all mutant measurements performed for these validation experiments (outlined below), we measured expression levels of 3 biological replicates of each mutant in a plate reader. The mutants, the *P*_*R*_ wildtype, and the negative control (the strain carrying the plasmid without a fluorescence marker in order to define ‘measurable expression’ – i.e. background fluorescence) were grown overnight in M9 minimal media supplemented with 0.2% CAS, 0.2% glucose, and 50μg/mL kanamycin. The overnight populations were diluted 1,000-fold, grown to OD_600_ of ∼0.1, and their fluorescence measured in a Bio-Tek Synergy H1 platereader. All the mutants were generated on the wildtype *P*_*R*_ plasmid backbone, using oligonucleotide cloning. We did not conduct tests for the variable spacer, because changes in spacer length in σ^70^-RNAP binding are well documented, so we assigned them their accurate biophysical meaning in the form of a binding energy penalty to altering spacer length away from the lowest energy spacer (9bp, corresponding to 17bp between -10 and -35 sites). We did not independently verify the clearance rate, as we use a simplified description of clearance rate that attempts to capture its effects in a single parameter, rather than explicitly accounting for the mechanisms of σ^70^-RNAP clearance.

#### Cumulative binding

To test for the role of multiple independent σ^70^-RNAP binding on gene expression levels, we created 29 specific promoters, which were derived from the *P*_*R*_ wildtype promoter. These promoters were selected so that the prediction of their gene expression levels would be different between two models: (i) the Standard model, which did not include cumulative binding, predicted no measurable expression for these promoters; (ii) a model that included only cumulative binding as a structural promoter feature, and which predicted measurable expression. We measured gene expression levels of these promoters in a plate reader as described above, compared the expression to the negative control through FDR-corrected t-tests, and found that the model with cumulative binding described experimental observations systematically better (Fig.S1).

We introduced additional mutations into a subset of these promoters in order to remove the additional σ^70^-RNAP binding sites. To do this, we used the Extended model to determine which mutations in the 29 promoters would reduce binding to any but the strongest binding site. Finding mutations that removed the secondary binding site(s) but that do not affect the binding to the strongest σ^70^-RNAP binding site was possible for only 7 of the 29 promoters. Removal of the additional binding site(s) in this manner led to a reduction of gene expression levels, and 6 of the 7 mutants exhibited no measurable expression – i.e. what was predicted by the Standard model which accounted for only a single binding site (Fig.S1).

#### Occlusive unproductive binding

We performed several different tests to validate various aspects of this structural feature. First, we created 20 promoters to compare predictions from two models: (i) model that allowed for cumulative binding, but all binding had a positive impact on expression; (ii) model that allowed for cumulative binding, but considered every binding that would not transcribe a complete ribosomal binding site as having a negative effect on expression. Model (i) predicted all 20 sequences to have measurable expression, while Model (ii), which accounted for occlusive unproductive binding, predicted no measurable expression for any of the sequences. By comparing the measured expression of each promoter to the negative control through FDR-corrected t-tests we found that only 2/20 mutants exhibited measurable expression, in agreement with the model that accounted for occlusive unproductive binding (Fig.S2A).

Second, we wanted to test if our model accurately predicted how expression levels change when an additional occlusive binding site is introduced. To this end, we started with a wildtype *P*_*R*_, and used the Extended model to generate a series of single point mutations that were predicted to gradually introduce a new binding site. σ^70^-RNAP binding to the newly introduced sites would result in transcripts without a complete ribosomal binding site. By constructing these mutants in the lab, we found that introduction of such a site into the *P*_*R*_ promoter indeed led to a significant reduction in expression levels (Fig.S2B). Note that the strongest σ^70^-RNAP binding in the wildtype *P*_*R*_ promoter is 28bp upstream of the ribosomal binding site.

We also did the reverse – starting with three promoters from the *P*_*R*_ mutant library that had a strong occlusive unproductive binding site, we used the Extended model to predict a series of single point mutants that would gradually remove this binding site. Experimental measurements of those mutants, when compared to the expression level of the starting promoter sequence through FDR-corrected t-tests, exhibited a significant increase in the expression levels as the binding site was removed – an increase that was consistent with that site acting in an occlusive manner, and not a cumulative one (Fig.S2C).

Finally, we wanted to determine the exact distance between the -10 element and the RBS that turned an additional binding site from being productive (cumulative) to being occlusive. To do this, we started with the same 3 promoters from the *P*_*R*_ mutant library that had a strong occlusive unproductive binding site as above, but this time we shifted the occlusive position of that binding site relative to the RBS. We shifted the binding site up to two positions closer to the RBS and up to seven positions further away from it, moving the binding site one position at a time - creating 9 mutants of each of the 3 starting promoters. We used FDR-corrected t-tests to compare the measured expression of each mutant to the original sequence they were mutated from. We found support for the hypothesis that, as the occlusive binding site was moved further away from the RBS, it became cumulative (Fig.S2D). This position matched the area determined with the model (see section *Towards the Extended model: occlusive unproductive binding*)

#### Occlusive binding on the reverse complement

In order to conduct an independent verification of this structural feature of promoters, we identified 4 promoters from the *36N* mutant library that the Extended model predicted to have strong occlusive binding on the reverse complement. We introduced up to 8 mutations into each of these 4 promoters that would progressively eliminate the occlusive site on the reverse complement. The introduced mutations had minimal predicted effect on binding on the productive strand (Fig.S3A). We measured gene expression levels of these mutants as described above, and performed a linear regression in order to correlate the measurements with the Extended model predictions of expression. For two sets of mutants, we found that the Extended model, which accounted for occlusive binding on the reverse complement, accurately predicted gene expression levels. For the other two mutant sets, removing the predicted binding site on the reverse complement had no measurable effect on expression levels (Fig.S3B). This data shows that occlusive binding on the reverse complement is a more complex promoter feature than what we accounted for in the Extended model. Nevertheless, including this promoter structural feature into the model led to a significant increase in predictability (especially of the *36N* dataset), which justified its inclusion into the Extended model.

### Verifying model predictions

Arguably the most surprising prediction arising from the Extended model is the ease of generating promoters from random non-expressing sequences. Specifically, we wanted to verify that single point mutations on random non-expressing 115bp-long sequences could generate dramatic shifts in expression levels. To do this, we experimentally created 20 pairs of promoters, each pair consisting of (i) a random non-expressing sequence; and (ii) the same sequence but with one point mutation that is predicted to lead to expression. We created these promoters on the *P*_*R*_ plasmid background, and measured gene expression levels as described in the section *Verifying promoter structural features*. By conducting a series of FDR-corrected, paired t-tests we found that, indeed, single point mutations improved gene expression levels for all but two of the 20 promoters (Fig.S7). Of the 20 original promoters, which were predicted not to have any expression, only two exhibited expression (Fig.S7), confirming the accuracy of the Extended model on sequences of 115bp length.

### Evolution simulations

We choose 100 random starting sequences of length 115bp, so that their predicted expression is less than measurable under both the Extended and the Standard model. We implement Gillespie-type simulation under the assumptions of Strong Selection and Weak Mutation, i.e. when we can assume no clonal interference. We define the time scale in the units of inverse mutation rate 1/*μ*, so in each iteration of the algorithm we simulate a single mutation that appears and gets fixed in the population with the probability given by the Kimura formula

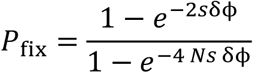

where *N* is the population size, δϕ is the change in the phenotypic value (gene expression level) due to mutation, and s is selection strength on the phenotype. Note that phenotype and selection are degenerate parameters: only their product represents the actual selection coefficient. We postulate ϕ = log_10_*P*_on_, so the typical mutational effects are in the range of 10^−3^ to 10^−2^. Under this regime, selection strength of 100 means a selection coefficient in the range 0.1—1.

We went through a sufficient number of iterations (new mutation events) until a threshold expression was reached by the population. We considered two thresholds: measurable expression we identified in our experiments to model the emergence of novel (weak) constitutive promoters; and the expression of the wildtype Lambda *P*_*R*_ promoter, as an example of a very strong promoter with high constitutive expression levels. For each of the 100 starting sequences, we performed 100 independent evolution runs, to obtain an estimate of the mean exit time (and the associated standard error of the mean) under the Standard and the Extended models.

We also varied the number of nucleotides in the 115bp-long sequence that were allowed to mutate. The mutagenized region was always in the center of the sequence, over which we always integrated to obtain expression. In simulations with a more constrained region allowed to mutate, the simulation would take too long until full convergence. To be robust against these cases, we stopped each simulation after 10N steps, truncating the distributions of evolution times for those sequences. Importantly, doing this introduced a bias that affected the Standard model more so than the Extended (because the Standard model took longer to reach the threshold), leading to an underestimate of how much faster the Extended model is compared to the Standard one. As such, what we report as the increase in the rates of evolution of the Extended model compared to the Standard one is likely the lower bound.

### Insights from applying the model to the E.coli genome

For all analyses described in this section, we used the *E.coli* K12 MG1655 genome (NCBI reference: NC_000913). Based on the genome annotations, we assigned one of the two identities for each of the nucleotide positions in that genome (Table S10): (i) ‘within genes’ (intragenic) — if the position was a part of the following annotation types: CDS or gene; (ii) Inter-genic — if none of the following annotation types were present at the position: misc_feature, mobile_element, repeat_region, tRNA, STS, tmRNA, rRNA, CDS, gene, ncRNA.

We calculated binding energy for each of the 5 x G configurations (5 spacer lengths, G is the number of positions/base-pairs in the genome), and then performed a thermodynamic sum over all spacer lengths for each position aligned so that each configuration coincided with the last base pair of the -10 element of the binding matrix. This way we obtained a free binding energy for each nucleotide (for each of the G positions) in the genome. We offset all the obtained free binding energy values so to set the minimum to zero, in the interest of later easier readability. Surprisingly perhaps, the nearest-neighbor correlation of the free energy values is negligible, so we did not subsample, but kept all values.

To assess whether there was selection against σ^70^-RNAP binding sites, we constructed a synthetic genome of 100 million base pairs with the same GC content as that region (inter-genic or within-genes) of the *E.coli* genome. We histogram the cumulative distributions, and normalize them to the number of base pairs (inter-genic or within-genes) relevant for the real *E. coli* sequence. For the p-value plot, we calculated the cumulative mass function of the Poisson distribution, at the value of the real histogram, and with mean given by the synthetic histogram value. This corresponded to the one-tailed p-value.

We also developed an alternative method of evaluating selection against σ^70^-RNAP binding sites, in order to strengthen the validity of our claims. This time, instead of creating a random synthetic genome and comparing the predicted expression levels across that genome using the σ^70^-RNAP energy matrix (Fig.1C), we created 100 shuffled energy matrices and evaluated free energies of such models across the actual *E.coli* genome. Matrices were permuted *per* position, meaning that the columns of the matrix were shuffled without altering the internal structure of each column. For each such matrix, we evaluated the model on every single position in the *E.coli* matrix and calculated cumulative histogram, as in the previous paragraph. The p-values were calculated assuming a normal distribution per bin. This is in fact a conservative estimate of the p-value, as for those bins with lower means, Gaussian overestimates the variance and hence the p-value. In the plot we illustrate the 95% confidence interval by showing explicitly the 3^rd^ and the 97^th^ percentile.

To predict the expression from known *E.coli* promoters, we assigned free energy to each of the 1,951 promoters in RegulonDB by integrating over a symmetric 40bp region around the reported transcript start. For a fair comparison in plot, we also integrated in the same way over 40bp chunks of all the intergenic regions described above (red line in Fig.3F). For each promoter, we then searched through RegulonDB for all transcription factors that bind it. If all of them were activators, we flagged that promoter as ‘activatable’, and similarly for ‘repressible’. If we found a mixture of activators and repressors affecting a given promoter, we flagged the promoter as ‘dual’. The promoters with no known transcription factor binding were flagged as ‘no info’.

## Notes

### Competing Interest Statement

The authors have declared no competing interest.

