## Supplementary material for "Structure and Evolution of Constitutive Bacterial Promoters": all supplementary figures and tables

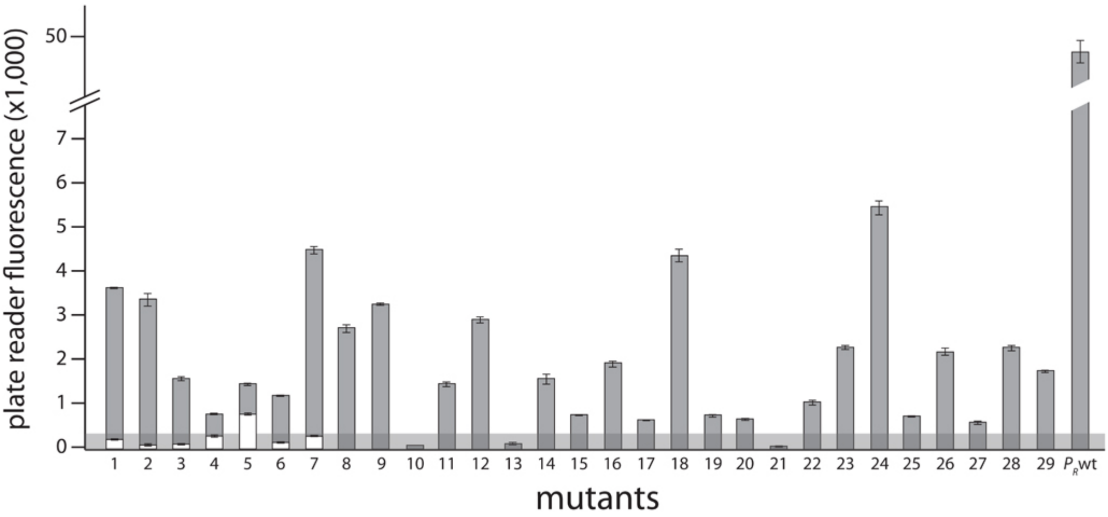


**Figure S1. Experimental verification of cumulative binding**. We experimentally created 29 promoters with the following property: the Standard model predicts no measurable expression from these promoters, while the Extended model predicts measurable expression due to the existence of multiple $\sigma^{70}$-RNAp binding sites. Fluorescence measurements are shown in grey bars, with error bars indicating standard error of the mean from 3 replicate biological measurements. The grey horizontal bar indicates the detectability ('no measurable expression') threshold. All promoters but 10, 13, and 21 exhibited significant measurable expression. We introduced additional mutations into promoters 1 to 7, in order to remove the secondary binding site(s) without affecting the strongest binding site. White bars show the expression levels of these additional mutants. Error bars are standard error of the mean from 3 replicate measurements. Only the mutated promoter 5 exhibited significant measurable expression.

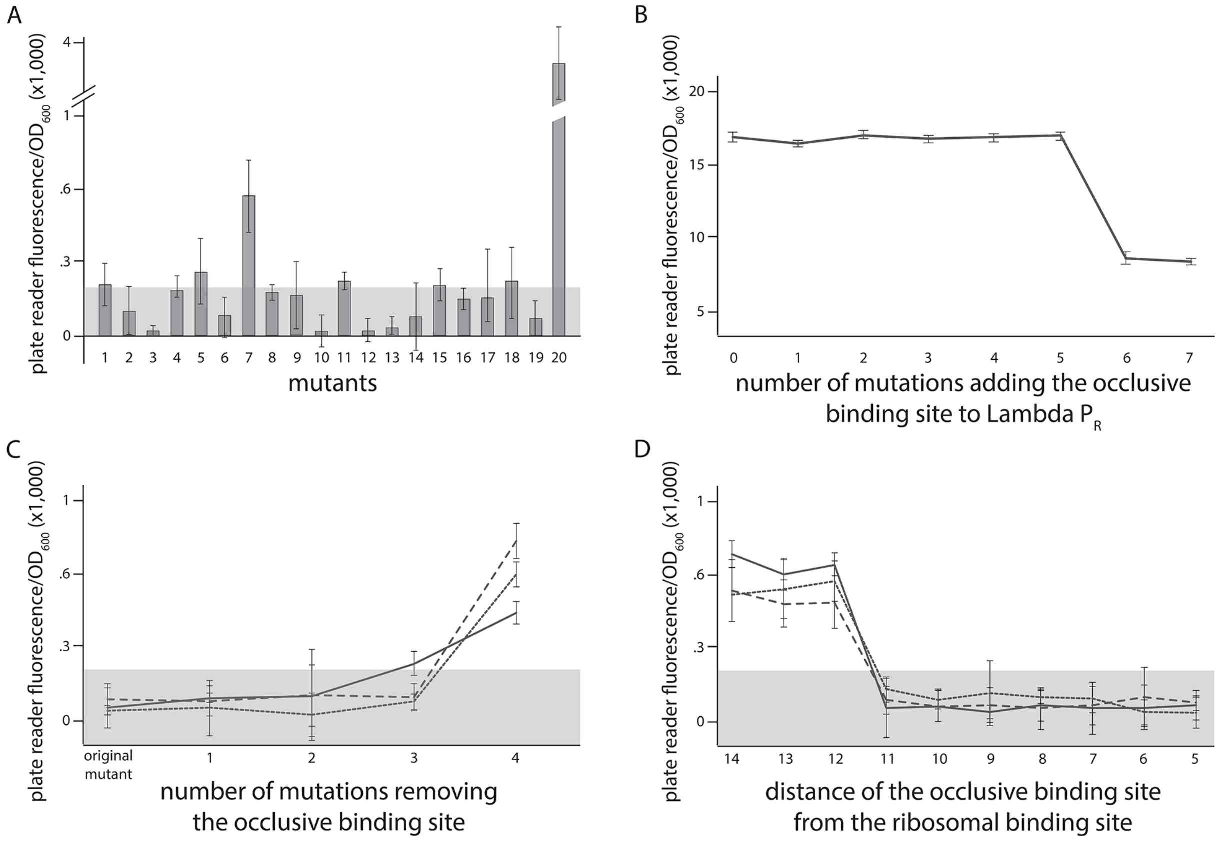


**Figure S2. Experimental verification of occlusive unproductive binding sites**. **A**) We created 20 promoter sequences for which the Extended model that accounts for occlusive unproductive binding predicted no measurable expression, while the model which did not account for occlusive unproductive binding predicted measurable expression. Bars are mean fluorescence measured from 3 biological replicates, and error bars are standard error of the mean. The grey shaded area indicates the detectability ('no measurable expression') threshold. Only mutants 7 and 20 exhibited significant measurable expression. **B**) We inserted mutations into the wildtype $P_R$ promoter to gradually introduce an additional binding site that was predicted to bind in an occlusive unproductive manner. These mutations were not predicted to significantly alter $\sigma^{70}$-RNAp binding to the existing dominant $P_R$ binding site. As mutations are introduced into the promoter, they generate stronger binding to the new site, which lowers gene expression levels. **C**) We mutated three promoters (originally found in the $P_R$ mutant library) to gradually remove their existing, predicted occlusive unproductive binding sites. The grey horizontal area indicates the detectability ('no measurable expression') threshold. As the predicted occlusive unproductive sites were removed, we measured a significant increase in gene expression levels. **D**) In order to experimentally verify the occlusive unproductive binding cut-off distance from the -10 end of the binding site to the beginning of the RBS, we started with the same three promoters as in **C**). We used the Extended model to identify the predicted occlusive unproductive binding site, and then we moved the site upstream and downstream to increase or decrease the distance from the RBS. We identified that a binding site that is 11 or fewer base pairs away from the RBS acts as an unproductive site, while those that are 12 or more base pairs away productively and cumulatively contributed to gene expression levels. This observation was in accordance with the cut-off identified by the model (Fig.S18).

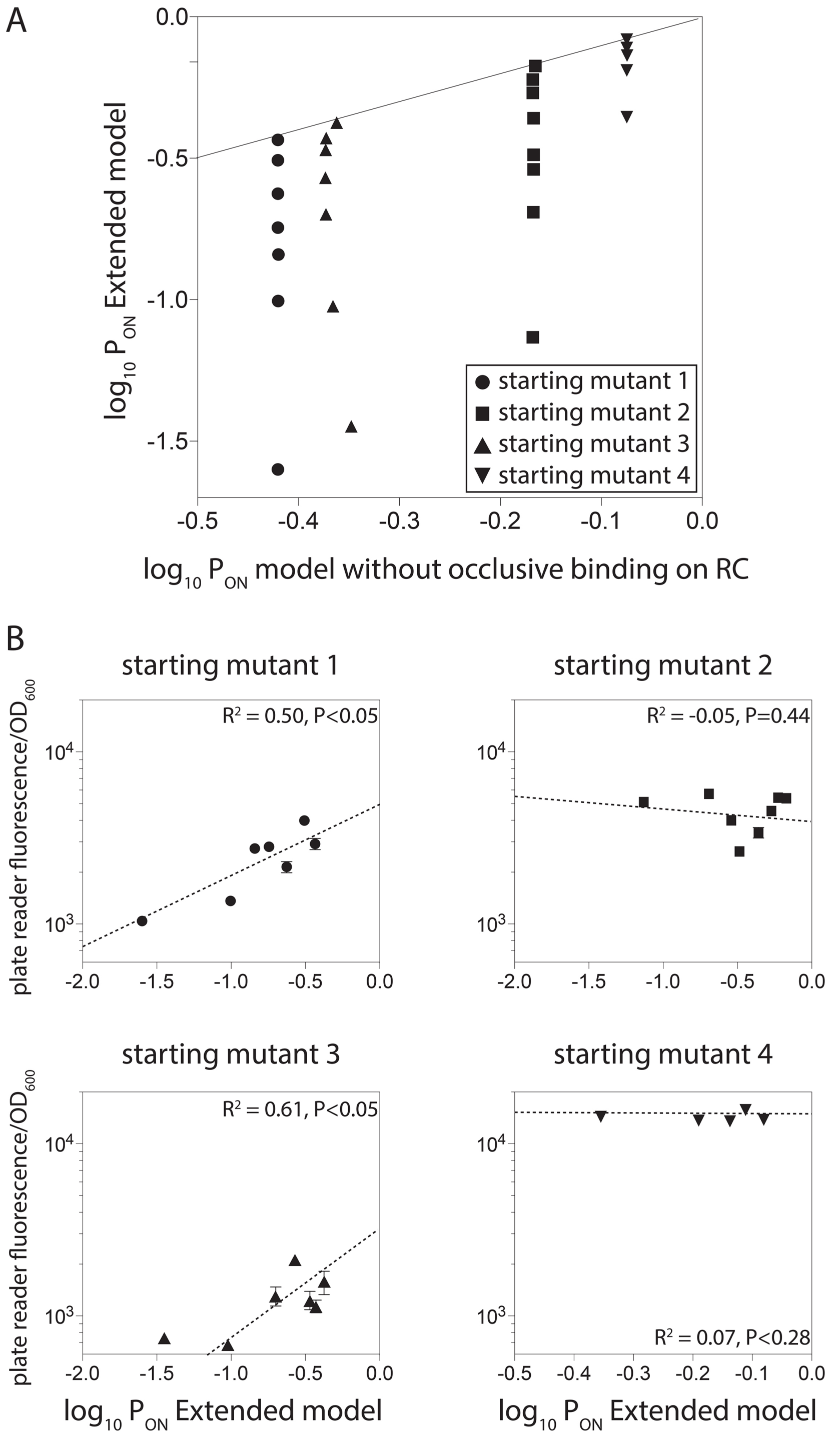


**Figure S3. Experimental verification of unproductive binding on the reverse complement**. **A**) We identified four promoter sequences for which we could introduce up to 8 mutations that would not alter predicted gene expression levels if the model did not account for unproductive binding on the reverse complement, but would if the Extended model was used. In other words, the 8 introduced mutations would gradually increase the strength of binding on the reverse complement while having a minimal effect on the strength of binding on the productive strand. **B**) Introduction of these mutations reduced the measured gene expression levels for two promoters (1 and 3) but had no effect on the expression levels from promoters 2 and 4.

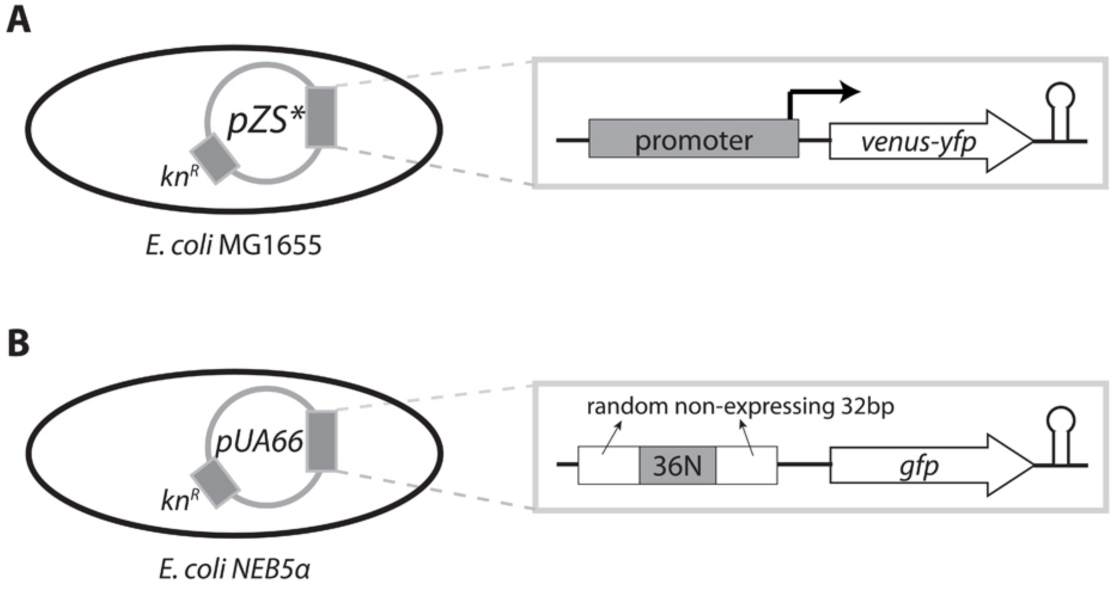


**Figure S4**. **Experimental plasmid systems**. **A**) For the $P_R$ and $P_L$ libraries, the synthetic construct used to detect the effects of promoter mutations consisted of a yellow fluorescent marker (*venus-yfp*), preceded by an RBS, and under the control of either the $P_R$ or $P_L$ promoter (or a $P_R$ or $P_L$ promoter mutant). The system was isolated from the rest of the plasmid by a T1 terminator (hairpin). This construct was placed on a small copy number pZS* plasmid (*SC101** origin) with kanamycin resistance, with *E.coli* MG1655 as host. **B**) The expression of a green fluorescence protein (*gfp*) was under the control of a random 100bp sequence consisting of: two 32bp-long random, non-expressing flanking sequences that were not mutated; and a 36bp-long sequence that was mutated randomly, with each nucleotide having 25% chance of being found at each position. This construct was placed on a pUA66 plasmid (*SC101* origin), with *E.coli* NEB5α as a host.

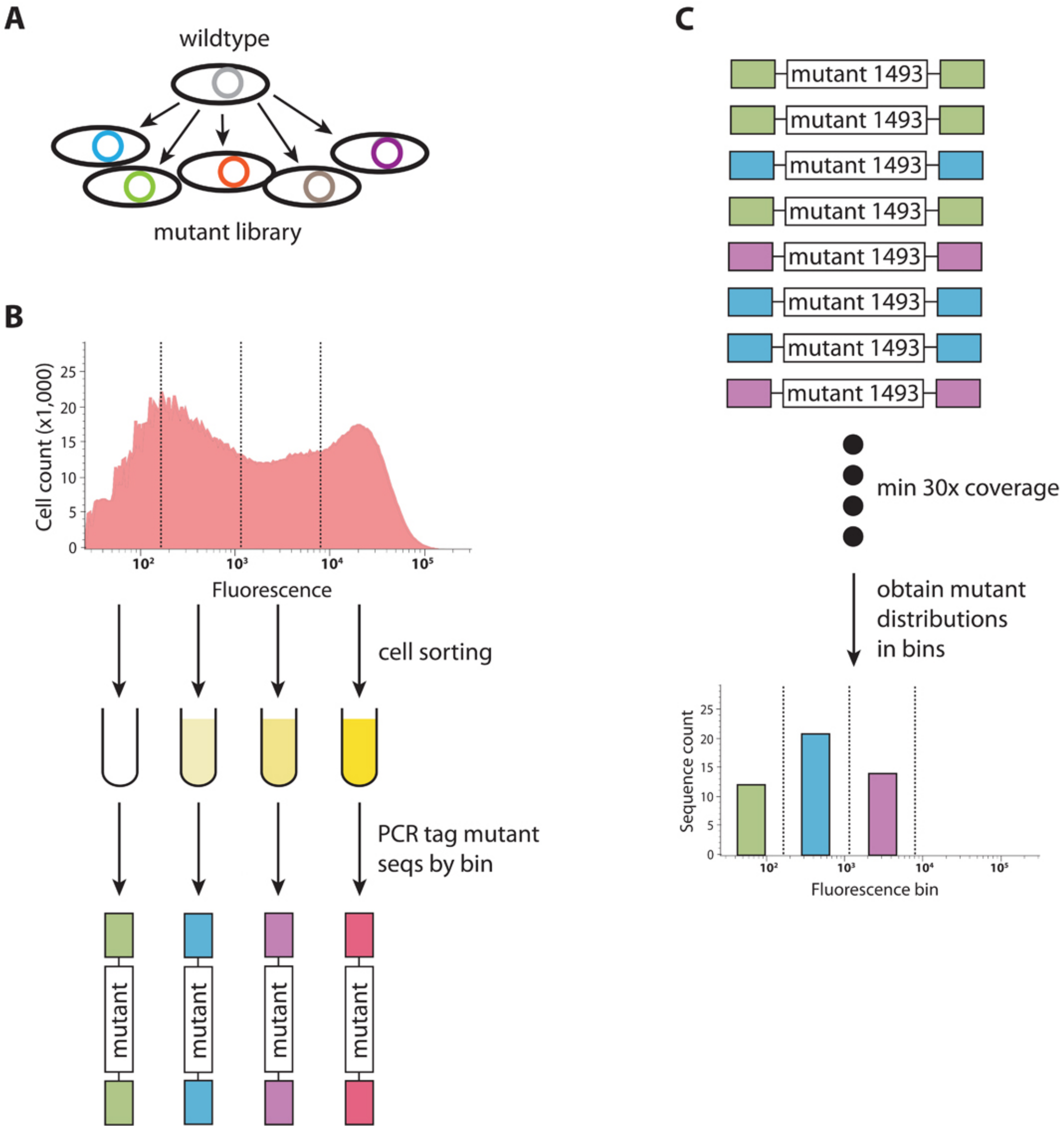


**Figure S5**. **Sort-seq experimental protocol**. **A**) Promoter mutants were cloned into the plasmid system using restriction/modification. The mutations were introduced at random, using pre-synthesized oligonucleotides with a fixed mutation rate (12% for the $P_R$, 9% for the $P_L$, and fully random for the 36N mutant library). The plasmids carrying mutant promoters were cloned either into MG1655 ($P_R$ and $P_L$ libraries) or NEB5α (36N library). **B**) Each random mutant library was sorted through Fluorescence Activated Cell Sorting (FACS) based on the fluorescence intensity detected at the single cell level. Mutants in $P_R$ and $P_L$ libraries were sorted into four, while the 36N library was sorted into 12 equidistant bins. 150bp-long fragments containing the promoter region of each sorted sub-library were PCR-tagged, and each library sequenced in bulk with 5 million total reads per library. **C**) We screened each sequence library for only those mutants that had at least 30x coverage, and obtained fluorescence distributions of each mutant across the bins.

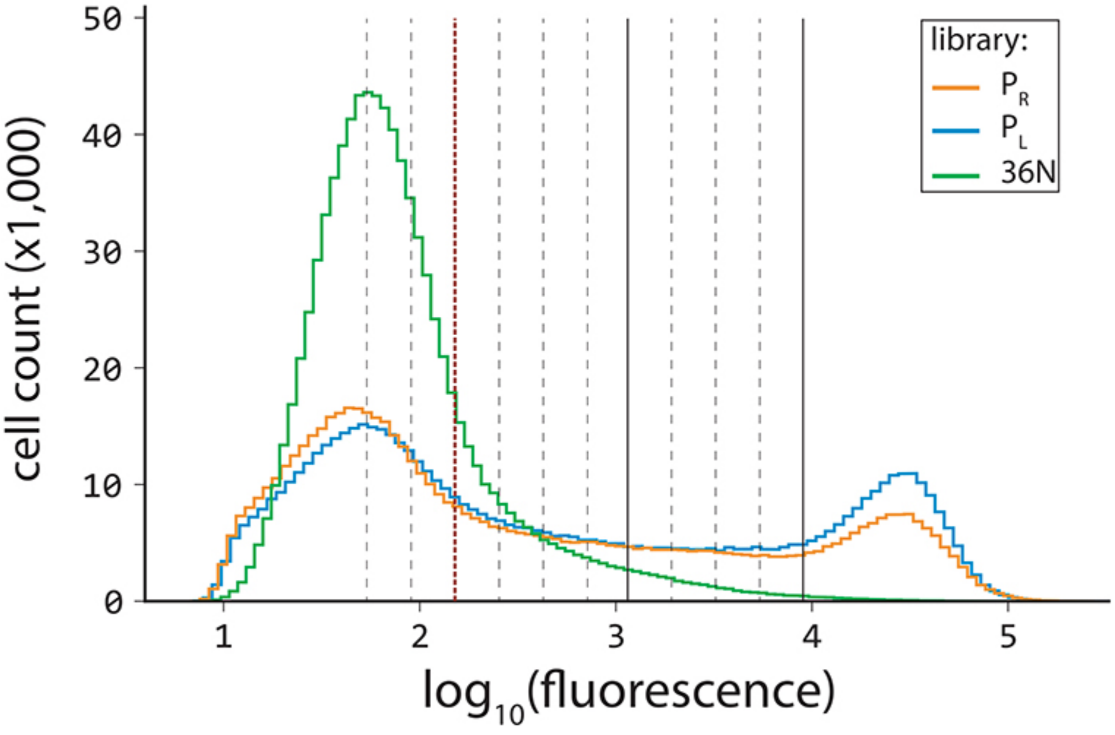


**Figure S6. Gene expression profiles for the three mutant libraries.** Flow cytometry measurements of one million mutants from each library showing distributions of fluorescence (as proxy for gene expression levels). The vertical red dotted line separates the mutants with no measurable expression (corresponding to Fig.3A). The red dotted line and the solid lines separate the four bins used to sort the $P_R$ and $P_L$ libraries (no, low, intermediate, and high expression, from left to right). The dotted lines mark the boundaries of the additional bins used to sort the *36N* library.

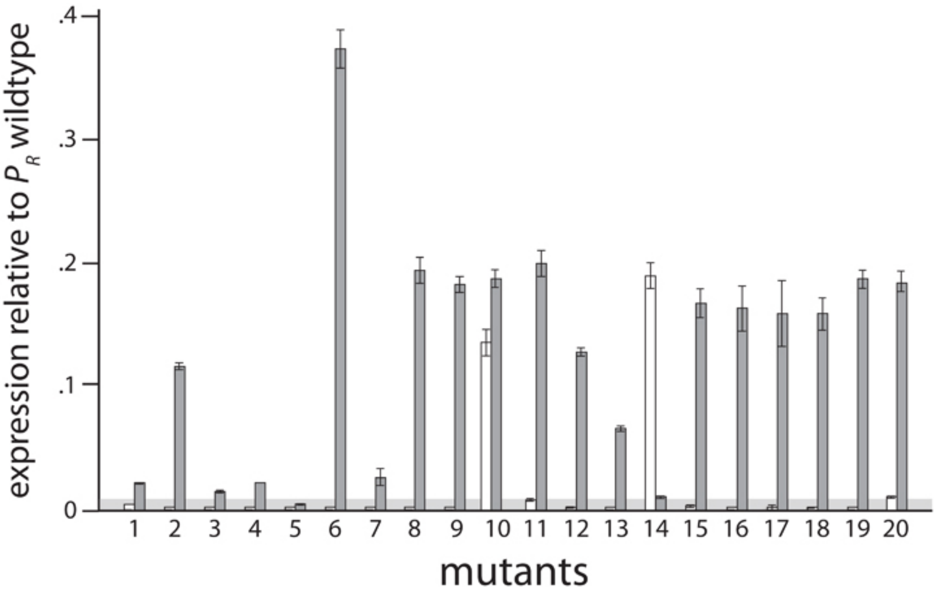


**Figure S7. Verifying model predictions on 115bp-long sequences**. In order to verify the ability of the Extended model to predict expression levels from 115bp-long promoters, and in particular to verify the model prediction of the ease of generating promoters from random non-expressing sequences, we generated 20 pairs of promoters. These pairs consisted of a randomly generated non-expressing sequence, and a sequence exactly one point mutation away that was predicted to have measurable expression. White bars are mean expression levels of three biological replicate measurements of non-expressing promoters; grey bars are the promoters with a single point mutation. Error bars are standard error of the mean. The grey horizontal bar indicates the detectability ('no measurable expression') threshold.

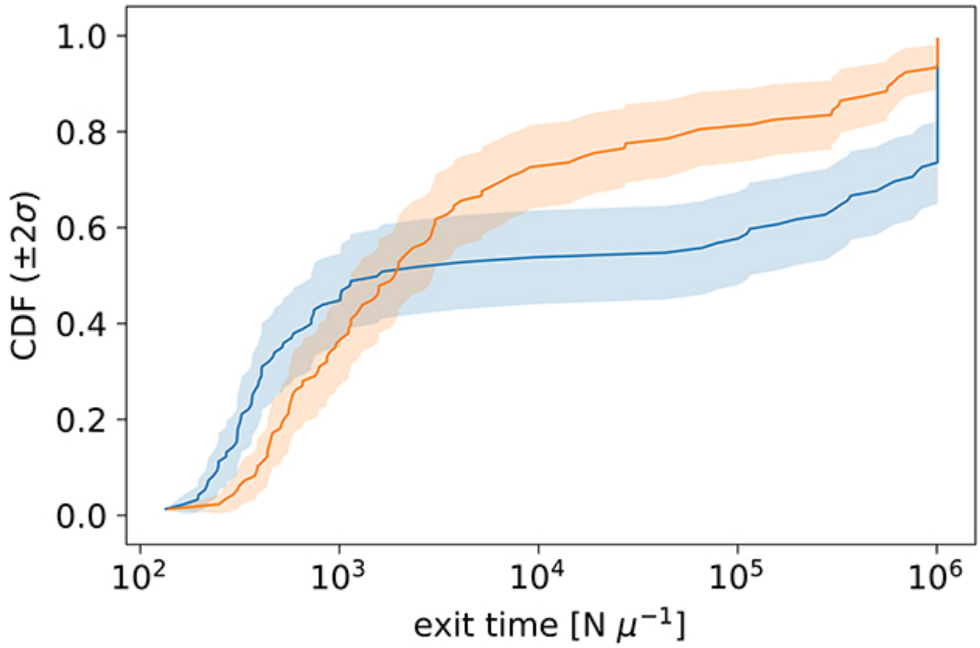


**Figure S8. Evolution times.** Cumulative distribution function (CDF) of the median times for promoter evolution under the Extended (orange) and Standard (blue) models for s=1, N=$10^4$, length of central mutagenized region of 70bp and high target expression level (that of wild type $P_R$). Evolution was simulated 100 independent times for each of the 100 starting random sequences. We present this specific set of parameters as this is the case where the largest fraction of simulations stopped at 10N iterations (our simulation limit), before reaching the target expression. For all parameter combinations, including the one shown here, more Standard model simulations terminate at 10N iterations compared to Extended model simulations. Taking a ratio of the mean time under this CDF for the Extended model over that for the Standard model therefore represents a conservative lower bound for the speedup in promoter evolution.

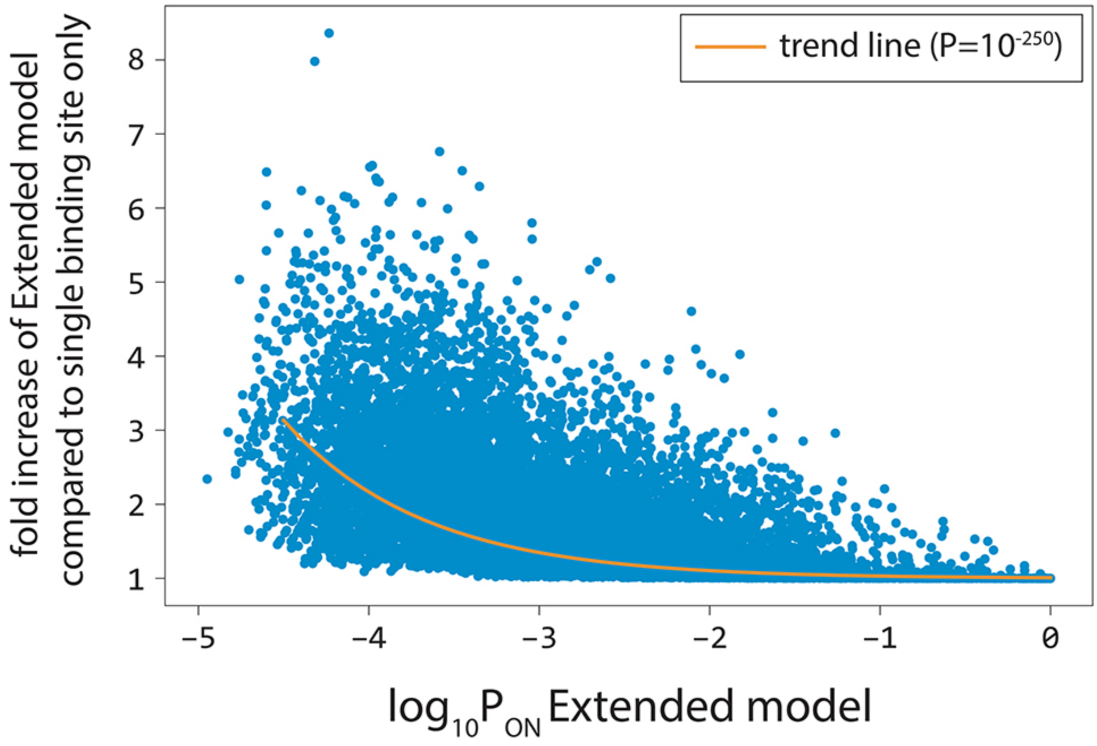


**Figure S9. Cumulative binding contributes more to expression at weak promoters**. For 100,000 random 100bp-long sequences, we calculated the fold increase in predicted gene expression levels of the Extended model compared to the model that is constrained to only the single strongest $\sigma^{70}$-RNAp binding site. Predicted expression levels from stronger promoters (higher $\log_{10}P_{on}$) were determined primarily by binding to the strongest $\sigma^{70}$-RNAp binding site. In contrast, predicted gene expression levels at weak promoters were more likely to be determined by $\sigma^{70}$-RNAp binding at multiple sites. The orange line is the trend line obtained through non-linear regression.

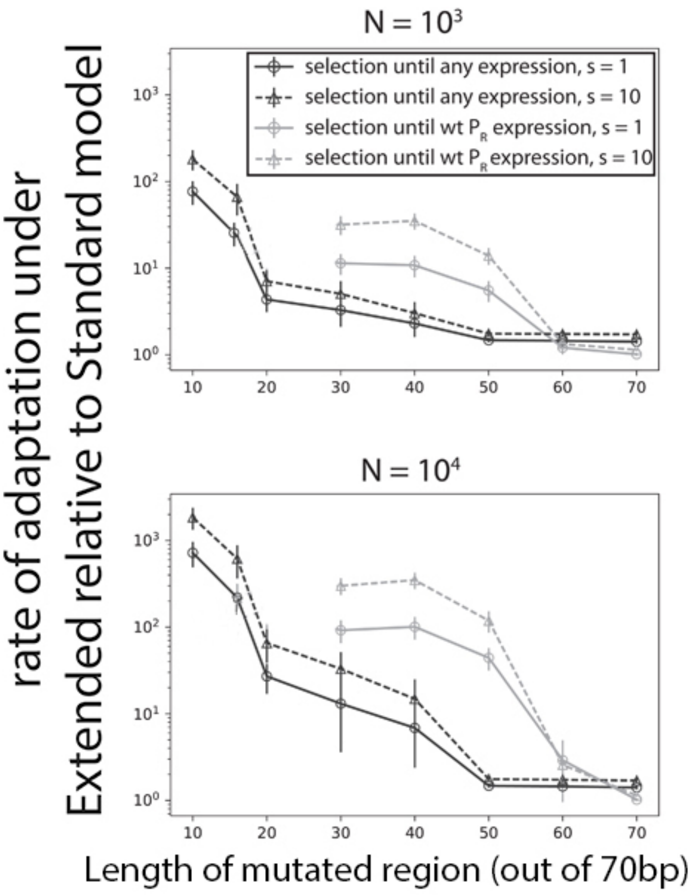


**Figure S10. Extended model predicts faster evolution under a range of conditions**. Selection at two different population sizes (top panel: $N=10^3$; bottom panel: $N=10^4$) using the Strong Selection Weak Mutation model at two selection strengths (s) and selecting to either $P_R$-levels of expression or any measurable expression. Selection was simulated through 100 independent runs for each of the 100 random starting sequences, with different lengths of the sequence allowed to mutate. Errors bars are standard errors of the mean across all replicates and starting sequences. Indicated selection refers to the selection on the phenotype difference ($\Delta\log_{10}P_{on}$).

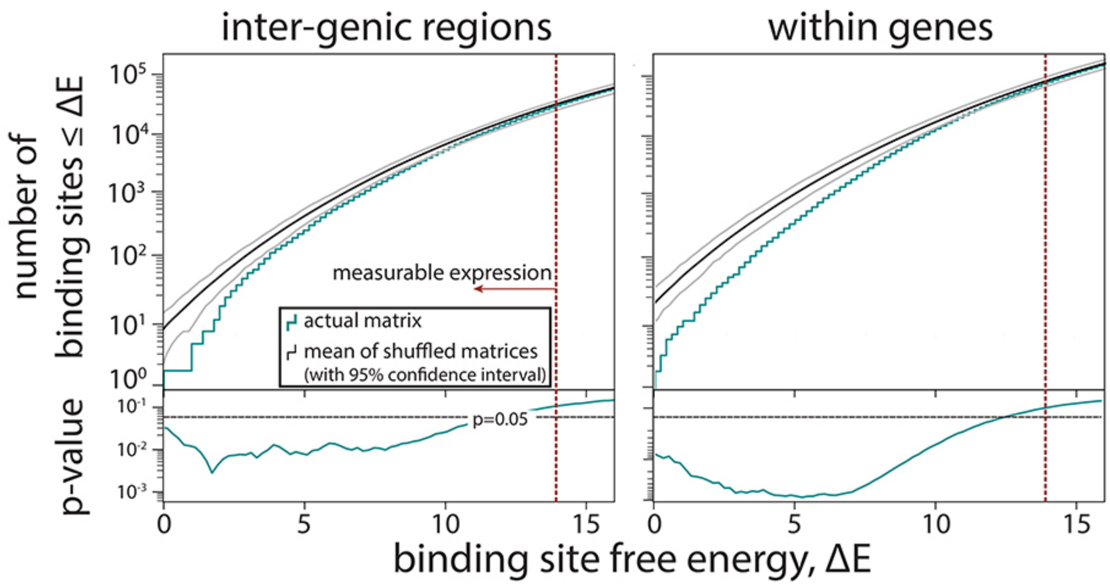


**Figure S11. Selection against σ$^{70}$-RNAP binding sites using shuffled binding matrices**. To provide an alternative measure to that presented in Fig.3E, instead of creating a random sequence and comparing the number of predicted σ$^{70}$-RNAP binding sites in it and in the *E.coli* genome, here we created 100 shuffled σ$^{70}$-RNAP energy matrices and used each of them to predict the expression from every single position in the *E.coli* genome. For each shuffle, we constructed cumulative histograms of free energy for inter-genic and within-genes regions. For each bin, we then calculated the p-value of the Extended model that used the actual σ$^{70}$-RNAP energy matrix, assuming a normal distribution with mean and standard deviation given by the set of models with shuffled matrices. This is a conservative estimate, as for energies $\Delta E<1$, the assumption of Gaussian distribution leads to overestimates of standard deviation. The matrices were shuffled per position, i.e., an energy matrix of dimension 4 x L, with L being the length of the binding site, is shuffled by randomly reordering the L columns while leaving the energy entries in each column unchanged in order and magnitude. Grey lines represent 95% confidence intervals.

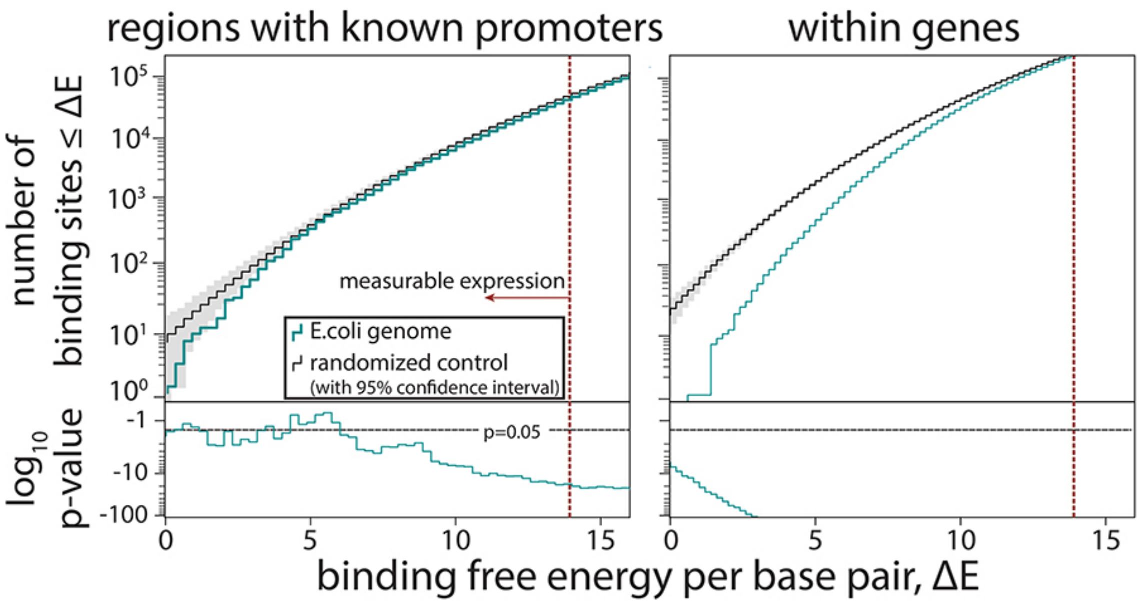


**Figure S12. Selection against $\sigma^{70}$-RNAp binding sites.** For evidence of selection against $\sigma^{70}$-RNAP binding sites only in the inter-genic regions that contain experimentally confirmed promoters (based on RegulonDB), we compared model-predicted binding energy across the region to the expected binding for a $10^8$bp random sequence with the GC% of the corresponding region. Also shown is the selection against binding sites within genes (same as in Fig.3E). Grey shaded areas are 95% confidence intervals.

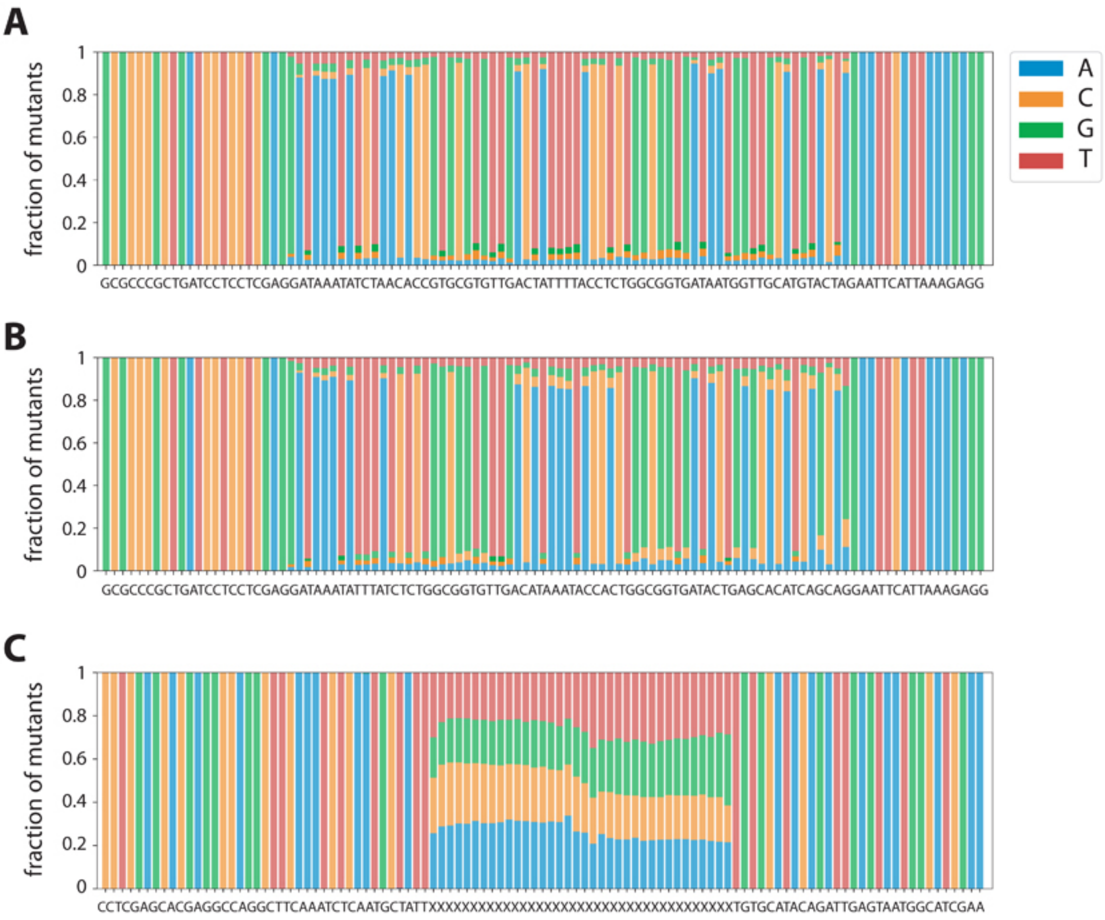


**Figure S13. Mutation frequencies in the three experimental libraries**. Fraction of mutants with a given nucleotide at each position is shown for **A**) $P_R$; **B**) $P_L$; and **C**) *36N* libraries. We did not observe any bias in the mutagenesis of libraries. The consensus sequence for each library is provided underneath each plot.

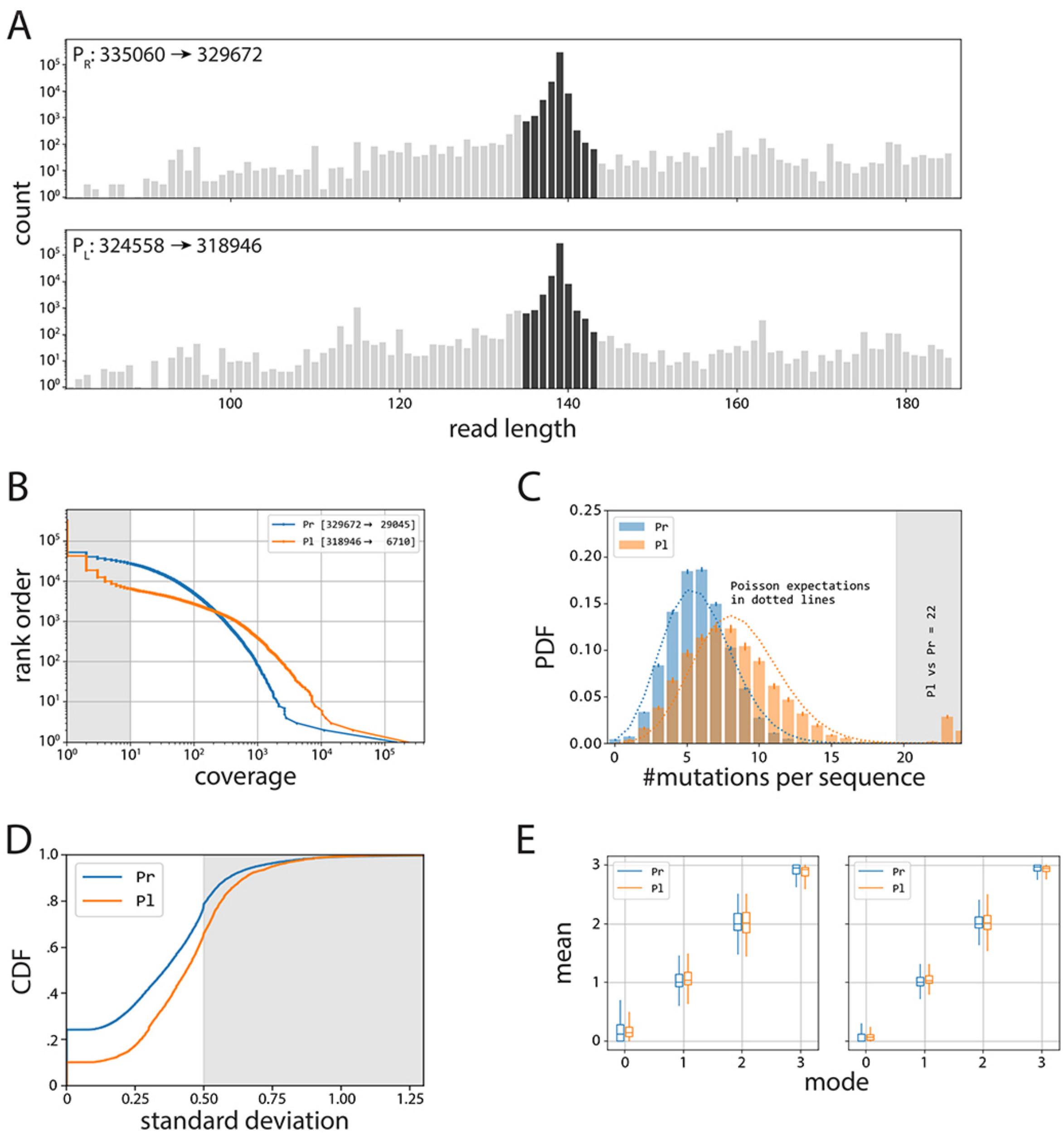


**Figure S14. Processing of $P_R$ and $P_L$ libraries**. **A**) All reads in the $P_R$ and $P_L$ libraries (grey), from which we take only those reads that are +/- 4bp away in length from the wildtype sequence (dark grey). **B**) Inverse cumulative distribution function (normalized to the total number of sequences), with shaded indicating the sequences we removed due to having less than 10x coverage. **C**) We removed sequences that had 20 or more single point mutations compared to their respective wildtype sequence. Note that this mainly affected the $P_L$ library (orange), as the original plasmid from which the libraries were cloned contained the wildtype $P_R$ sequence. **D**) Cumulative distribution function (CDF) of standard deviation of expression bin numbers, with shaded sequences the ones we removed from subsequent analyses. **E**) Box plots indicating the distributions of mean values (in bin units) for a given mode (in bin units)**,** before (left) and after (right) selecting for only those where mean, mode and median are within 0.5.

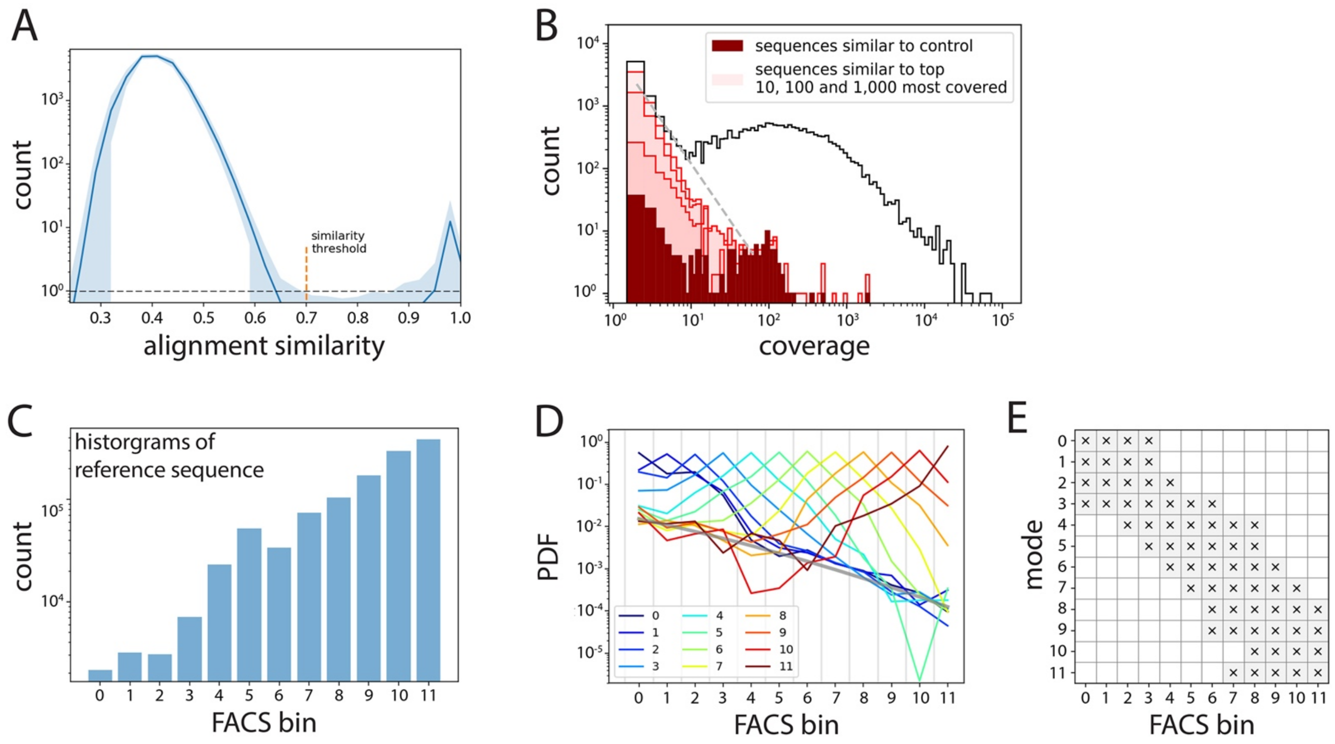


**Figure S15. Processing of the 36N library**. **A**) Average histogram of alignment similarity for the 1000 most covered sequences (shaded area indicates 95% confidence interval). We used the similarity threshold of 0.7 between low- and high-scoring modes to select for unique sequences and eliminate sequencing errors. **B**) Histogram of coverage (black line), with highlighted contributions of the noise cloud around the reference sequence (dark red), and the clouds around the 10, 100 and 1000 most abundant sequences (from darkest to lightest shade of red, respectively). **C**) Histogram of counts for the reference sequence per bin, used to debias all other distributions. **D**) Template probability distribution functions obtained as averages of PDFs that have the same mode (indicated by color). The inferred FACS noise background is shown as a thick grey line. Given a distribution, we only accepted values in the bins in which the appropriate reference was three times above the inferred background. Such filter is shown in **E**)

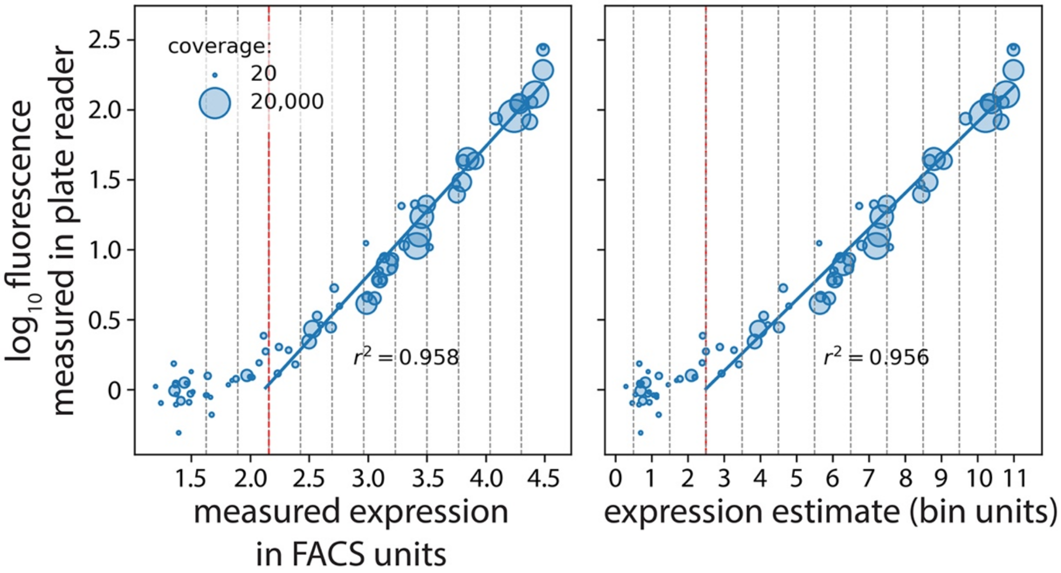


**Figure S16**. **Plate reader validation of 36N data processing**. 77 mutants (with an approximately equal number of mutants selected from each of the 12 bins) were selected randomly and their expression levels measured in a plate reader. We correlated their expression measured in the plate reader with our estimates in FACS units (left) and bin units (right). The vertical red dotted line marks the measurable expression threshold in the flow cytometer. Measured expression in FACS units and expression estimate are shown in log scale.

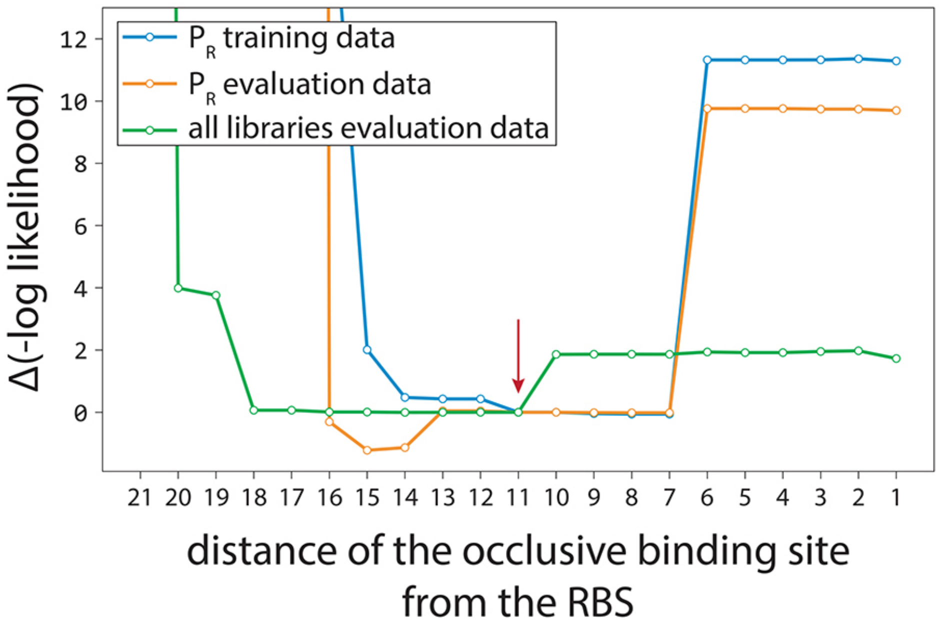


**Figure S17. Model determination of occlusive unproductive binding sites**. We evaluated the Extended model fitted on the $P_R$ training dataset, with varying thresholds between productive and occlusive unproductive bindings. Shown is the change in the negative log likelihood on the dataset indicated in the legend. Red arrow indicates the actual threshold ultimately used in the model. Because this modeling only provided a range for the productive/unproductive cut-off distance between the RNAP binding site and the RBS, we carried out dedicated experiments to systematically validate and probe occlusive unproductive binding (Fig.S2D).

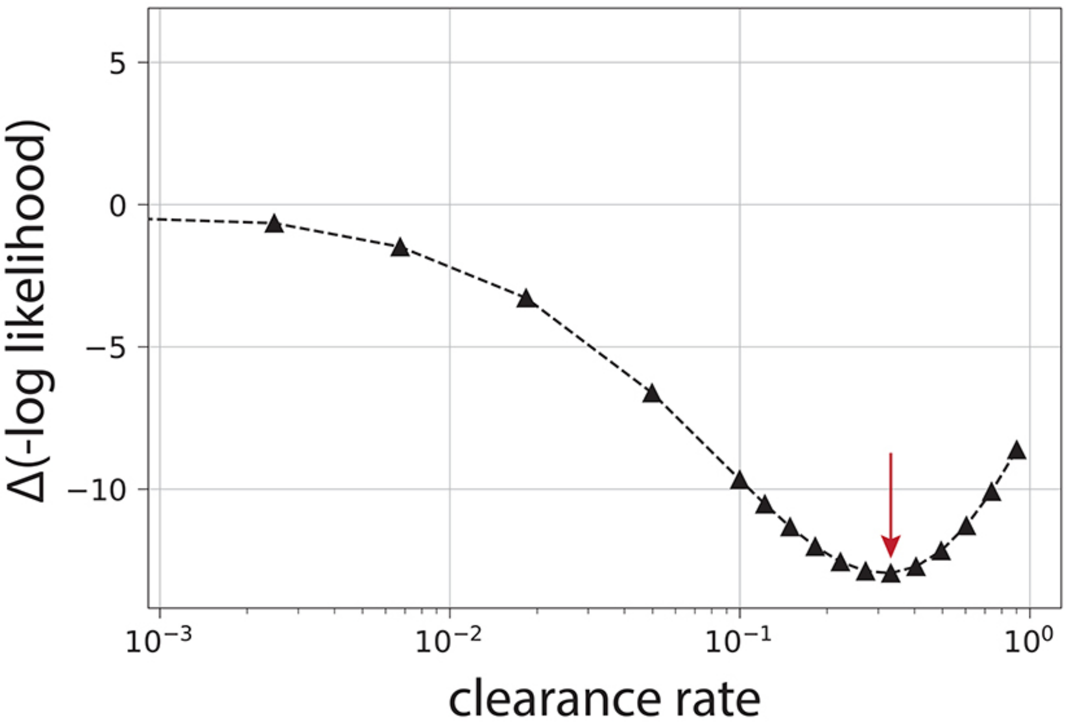


**Figure S18**. **Optimal value of the clearance rate**. We scanned through the possible values of the relative clearance rate of the $\sigma^{70}$-RNAp complex from the promoter, using the Extended model fitted on the training subsets of all mutant libraries. For each value, we refit chemical potential and hyperparameters of logistic regression using the training dataset. Shown is the change in negative log likelihood on the training data. The optimal value is indicated with the red arrow, though a wider range of values is compatible with the data. For the majority of values in the compatible range including the optimal value, the model performance improves also on the validation dataset.

**Table S1. Inferred values for model parameters**. For each position in the energy matrix, the energy penalty is normalized to the lowest energy (strongest binding) residue in that position, which is set to zero. Same applies for the spacer variation penalty, where the optimal spacer length is set to zero. Orange background marks the canonical -10 and -35 binding sites of $\sigma^{70}$-RNAP.

| | Residue | A | C | G | T |
|---|---|---|---|---|---|
| | most upstream position | 1.0042 | 0.8725 | 0.9656 | 0 |
| | | 0 | 0.7132 | 0.2516 | 1.4211 |
| | | 1.4277 | 1.5981 | 0.9806 | 0 |
| | | 0.8669 | 0.9585 | 0 | 0.4791 |
| | | 0.673 | 0.3131 | 0 | 0.3544 |
| **Energy Matrix -10 foot** | | 4.8851 | 3.7094 | 4.566 | 0 |
| | | 0 | 5.8406 | 6.3724 | 4.2602 |
| | | 1.1993 | 1.3188 | 1.5041 | 0 |
| | | 0 | 1.9175 | 2.0824 | 1.7399 |
| | | 0 | 0.8898 | 2.1099 | 2.3318 |
| | | 4.2699 | 4.646 | 4.8871 | 0 |
| | most downstream position | 0 | 0.8024 | 0.3061 | 0.3178 |

| | | | | | |
|---|---|---|---|---|---|
| | most upstream | 0 | 0.6782 | 0.2726 | 0.1205 |
| | | 0.3426 | 0 | 0.2543 | 2.0091 |
| | | 4.1376 | 4.9489 | 3.7397 | 0 |
| | | 2.2569 | 3.995 | 2.4141 | 0 |
| | | 5.4228 | 4.5391 | 0 | 2.2583 |
| **Energy Matrix -35 foot** | | 0 | 1.621 | 3.9457 | 2.3814 |
| | | 1.7288 | 0 | 2.1718 | 1.5128 |
| | | 0 | 1.3907 | 1.1822 | 0.5051 |
| | | 0.4337 | 0.7953 | 0.9523 | 0 |
| | | 0.0977 | 0.4765 | 0.5841 | 0 |
| | | 0.2763 | 0.1117 | 0.4113 | 0 |
| | most downstream | 0.0844 | 0.1982 | 0 | 0.0121 |

| | | | | | | |
|---|---|---|---|---|---|---|
| **Spacer** | difference to optimal (9bp) spacer length | -2 | -1 | 0 | 1 | 2 |
| | energy penalty | 8.514 | 2.122 | 0 | 1.242 | 5.346 |

| | | |
|---|---|---|
| **Clearance rate** | | 0.332 |

| | | | |
|---|---|---|---|
| **Chemical potential** | library | *$P_R$ / $P_L$* | *36N* |
| | | 11.128 | 10.531 |

**Table S2. All significant pairwise (dinucleotide) interactions.** 'Identity' refers to the two positions and specific residues that have a significant positive (destabilizing) or negative (stabilizing) interaction, with numbers indicating the position of the residue in the matrix shown in Fig.1C. Shaded cells indicate the strongest interactions, which are shown in Fig.1C. The numbers indicate the position in the energy matrix, starting from the most upstream to the most downstream (left to right in the energy matrix shown in Fig.1C).

| Identity | Interaction strength |
|---|---|
| (2G, 24A) | -0.076 |
| (9A, 10A) | -0.083 |
| (8T, 9T) | -0.091 |
| (23T, 24G) | -0.092 |
| (22T, 26G) | -0.094 |
| (24A, 25T) | -0.094 |
| (24A, 26G) | -0.101 |
| (27T, 28A) | -0.212 |
| (6C, 7A) | -0.238 |
| (5C, 6C) | -0.275 |
| (6T, 7G) | -0.308 |
| (4C, 9C) | -0.342 |
| (6A, 7A) | 0.058 |
| (23G, 29C) | 0.062 |
| (6A, 32A) | 0.092 |
| (25G, 26G) | 0.106 |
| (6C, 7C) | 0.109 |
| (23T, 25T) | 0.123 |
| (23T, 26G) | 0.127 |
| (2A, 24A) | 0.128 |
| (24A, 26T) | 0.135 |
| (23A, 28G) | 0.150 |
| (6A, 7G) | 0.155 |
| (22A, 23A) | 0.168 |
| (22A, 24C) | 0.170 |
| (2A, 27T) | 0.179 |
| (23T, 24C) | 0.188 |
| (1G, 27T) | 0.243 |
| (25A, 27T) | 0.257 |
| (23A, 27T) | 0.329 |
| (23T, 24T) | 0.390 |

**Table S3. Improvement to predictability based on each promoter feature**. Each structural promoter feature was added to the simpler iteration of the model, starting from the Standard model and building progressively towards the Extended. The clearance rate and dinucleotide interactions were included only in the model fitted on all three libraries, and not on the model iterations fitted only on the $P_R$ library. The values are the fraction of variance explained of the evaluation dataset.

| | | ***Library used for evaluation*** | | |
|---|---|---|---|---|
| **Model** | **Library used for fitting** | $P_R$ | $P_L$ | ***36N*** |
| **Standard** | $P_R$ | 0.7467 | 0.6480 | 0.5098 |
| **+flexible spacer** | $P_R$ | 0.7989 | 0.6954 | 0.6476 |
| **+cumulative binding** | $P_R$ | 0.8089 | 0.7053 | 0.6549 |
| **+occlusive binding = Extended** | $P_R$ | 0.8091 | 0.7042 | 0.6574 |
| **Standard** | all three libraries | 0.7076 | 0.6255 | 0.6173 |
| **+flexible spacer** | all three libraries | 0.7838 | 0.6925 | 0.7682 |
| **+cumulative binding** | all three libraries | 0.7928 | 0.7085 | 0.7728 |
| **+occlusive binding** | all three libraries | 0.7865 | 0.7068 | 0.7754 |
| **+clearance rate** | all three libraries | 0.7872 | 0.7070 | 0.7757 |
| **+dinucleotide interactions** | all three libraries | 0.7932 | 0.7132 | 0.7893 |

**Table S4. Processing of the $P_R$ and $P_L$ mutant libraries**. The table shows the number of reads remaining in the datasets following each step of data processing, from original sequenced library down to the final library used for model fitting and evaluation.

| | $P_R$ | $P_L$ |
|---|---|---|
| **Initial number of reads** | 7,138,685 | 9,149,460 |
| **Filtered on 0 mismatches** | 5,432,101 | 6,557,151 |
| **Condition on same (and valid) left and right bin tags** | 2,637,166 | 2,459,553 |
| **Number of unique sequences** | 335,060 | 324,558 |
| **Condition on length being within 4bp of canonical** | 329,672 | 318,946 |
| **Condition on coverage of at least 10** | 29,045 | 6,710 |
| **Cond. on Shine-Dalgarno within +/- 5bp of canonical** | 29,031 | 6,694 |
| **Remove sequences too different from ancestor** | 29,020 | 6,415 |
| **Remove sequences with expression st.dev. > 0.5** | 22,884 | 4,239 |
| **Condition on median, mode and mean of expression distribution being within 0.5** | 22,769 | 4,222 |
| **Condition on coverage of at least 30** | 12,476 | 2,984 |

**Table S5. Processing of the *36N* mutant library**. The table shows the number of reads remaining in the dataset following each step of data processing, from original sequenced library down to the final library used for model fitting and evaluation.

| | ***36N*** |
|---|---|
| **Initial number of reads** | 10,124,219 |
| **Filtered on 0 mismatches** | 9,917,488 |
| **Discard (yet save) reads that map to control sequence** | 8,772,436 |
| **Discard reads that cannot map left and right flanking region** | 7,031,460 |
| **Condition on the length of the core region within 2bp of canonical** | 6,498,273 |
| **Number of unique sequences** | 90,071 |
| **Condition on coverage of at least 2 (temporarily)** | 24,527 |
| **Condition on coverage of at least 30** | 13,341 |

**Table S6. Sizes of datasets after the splits**

| | Training | Validation | Evaluation | Total |
|---|---|---|---|---|
| $P_R$ | 7,485 | 2,495 | 2,496 | 12,476 |
| $P_L$ | 1,790 | 597 | 597 | 2,984 |
| ***36N*** | 8,004 | 2,668 | 2,669 | 13,341 |

**Table S7. Number of mutants per expression bin for each split of the $P_R$ dataset**. Bins are no ('0'), low ('1'), intermediate ('2') and high ('3').

| $P_R$ | **0** | **1** | **2** | **3** |
|---|---|---|---|---|
| **evaluation** | 276 | 847 | 450 | 923 |
| **validation** | 290 | 848 | 437 | 920 |
| **training** | 831 | 2,363 | 1,375 | 2,916 |

**Table S8. Number of mutants per expression bin for each split of the $P_L$ dataset**. Bins are no ('0'), low ('1'), intermediate ('2') and high ('3').

| $P_L$ | **0** | **1** | **2** | **3** |
|---|---|---|---|---|
| **evaluation** | 152 | 144 | 88 | 213 |
| **validation** | 173 | 130 | 90 | 204 |
| **training** | 450 | 399 | 246 | 695 |

**Table S9. Number of mutants per expression bin for each split of the 36N dataset**. Bins are ordered from lowest ('0') to highest ('11').

| ***36N*** | **0** | **1** | **2** | **3** | **4** | **5** | **6** | **7** | **8** | **9** | **10** | **11** |
|---|---|---|---|---|---|---|---|---|---|---|---|---|
| **evaluation** | 177 | 1,477 | 424 | 220 | 107 | 77 | 87 | 45 | 26 | 16 | 7 | 6 |
| **validation** | 164 | 1,509 | 453 | 226 | 105 | 65 | 70 | 42 | 15 | 11 | 7 | 1 |
| **training** | 483 | 4,535 | 1,296 | 648 | 345 | 216 | 205 | 124 | 72 | 33 | 22 | 25 |

**Table S10. *E.coli* genome partitioned into within genes and inter-genic regions**.

| **Regions** | **number of bp** | **percent of the genome** | **GC content** |
|---|---|---|---|
| **whole genome** | 4,641,652 | 100 % | 50.8 % |
| **within genes** | 4,158,349 | 89.6 % | 51.8 % |
| **inter-genic** | 446,896 | 9.6 % | 41.1 % |
